# FDA-Approved Drug Screening in Patient-Derived Organoids Demonstrates Potential of Drug Repurposing for Rare Cystic Fibrosis Genotypes

**DOI:** 10.1101/2022.09.02.506304

**Authors:** E. de Poel, S. Spelier, M.C. Hagemeijer, P. van Mourik, S.W.F. Suen, A.M. Vonk, J.E. Brunsveld, G. N. Ithakisiou, E. Kruisselbrink, H. Oppelaar, G. Berkers, K.M. de Winter-de Groot, S. Heida-Michel, S.R. Jans, H. van Panhuis, M. Bakker, R. van der Meer, J. Roukema, E. Dompeling, E.J.M. Weersink, G.H. Koppelman, A.R. Blaazer, J.E. Muijlwijk-Koezen, C.K. van der Ent, J.M. Beekman

## Abstract

**Background:** Preclinical cell-based assays that recapitulate human disease play an important role in drug repurposing. We previously developed a functional forskolin induced swelling (FIS) assay using patient-derived intestinal organoids (PDIOs), allowing functional characterization of CFTR, the gene mutated in people with cystic fibrosis (pwCF). CFTR function-increasing pharmacotherapies have revolutionized treatment for approximately 85% of people with CF, but a large unmet need remains to identify new treatments for all pwCF.

**Methods:** We used 76 non-homozygous F508del-CFTR PDIOs to test the efficacy of 1400 FDA-approved drugs on improving CFTR function, as measured in FIS assays.

**Results:** Based on the results of a secondary validation screen, we investigated CFTR elevating function of PDE4 inhibitors and currently existing CFTR modulators in further detail. We show that PDE4 inhibitors are potent CFTR function inducers in PDIOs and that CFTR modulator treatment rescues of CF genotypes that are currently not eligible for this therapy.

**Conclusions:** This study exemplifies the feasibility of high-throughput compound screening using PDIOs and we show the potential of repurposing drugs for pwCF that are currently not eligible for therapies.

**One-sentence Summary:** We screened 1400 FDA-approved drugs in CF patient-derived intestinal organoids using the previously established functional FIS assay, and show the potential of repurposing PDE4 inhibitors and CFTR modulators for rare CF genotypes.

## INTRODUCTION

Preclinical cell-based assays that recapitulate human disease play an important role in the first steps of drug development. Drug repurposing is the process of using clinically approved drugs outside their original disease-indication ^1^. Pharmacokinetic and safety data that is readily available for existing drugs can enable a rapid use in clinical studies, which is particularly relevant in the context of rare diseases and personalized medicine where small patient populations enlarge economic and technical complexities. It has been estimated that 75% of known drugs could potentially be repositioned for various diseases ^2^.

Cystic fibrosis (CF) is a rare hereditary disease caused by mutations in the *CFTR* gene. Pharmacotherapies termed CFTR modulators that rescue CFTR function have revolutionized treatment for approximately 85% of people with CF (pwCF) who carry the most prevalent F508del-CFTR mutation ^3^, but a large unmet need remains to identify new and affordable treatments for patients with CFTR mutations that are non-eligible or non-responsive for CFTR modulators. Such mutations encompass several classes, ranging from Class I mutations, such as premature termination codon (PTC) and splicing mutations, to very rare uncategorized mutations. CFTR function measurements in patient-derived intestinal organoids (PDIO’s) associate with clinical features of CF and may enable drug repurposing in a personalized setting ^4^. These CFTR function measurement are performed by means of the forskolin-induced (FIS) assay, in which forskolin induces fluid secretion into the organoid lumen resulting in rapid organoid swelling in a (near-to) complete CFTR-dependent manner ^5,6^. As found by us and others, CFTR function measurements in PDIOs associate with disease severity indicators of CF and CFTR modulator response, thereby enabling drug discovery efforts^7,8^. The established correlation between the FIS assay and clinical response furthermore allows theratyping, or the matching of patients to beneficial compounds based on laboratory results of the patient-derived cells. The fact that the FIS assay is well characterized in regards to translation of results to the pwCF in the clinic, indicates its potential for drug repurposing experiments.

Other prerequisites for drug repurposing are that the exploited assay is high-throughput and robust. We recently succeeded in establishing a high-throughput screening version of the FIS assay, allowing testing of compounds that directly or indirectly influence CFTR function in a high-throughput manner on CF patient-derived material ^9^. Using this miniaturized FIS assay, we screened 76 non-homozygous F508del PDIOs, aiming to identify CFTR function enhancing drugs in a 1400-compound FDA-approved drug library. The included PDIOs represent those

CF patients who are not eligible for therapies at present-day. Three main hit families were distinguished: existing CFTR modulators, PDE4 inhibitors and tyrosine kinase inhibitors (TKIs). Due to the toxic nature of the latter category and the fact that PDE4 inhibitors are already used for treatment of the airway disease COPD ^10^, we argued that repurposing of PDE4 inhibitors and extending the label of CFTR modulators for currently non-approved genotypes, hold most potential. We investigated those families in the rest of this study in several ways, amongst which characterization of PDE expression, characterization of responsive PDIOs and assessing the correlation between PDIO response to CFTR modulators and clinical efficacy of those modulators.

This study exemplifies the feasibility of high-throughput compound screening using PDIOs in an assay with a functional read-out. We show the potential of repurposing drugs for people with CF carrying non-F508del genotypes that are currently not eligible for therapies, underlining the need for label expansion of CFTR modulators for currently non-eligible pwCF. Additionally, we describe the potential therapeutic benefit of PDE4 inhibitors for pwCF with residual functional CFTR mutations. Altogether, this study underlines the potential and importance of identification of potential treatments and responsive patients, paving the way for patient stratification in the upcoming era of personalized medicine.

## RESULTS

### FDA-approved Drugs Increase FIS in Non-Homozygous F508del PDIOs

We set out to study rescue of CFTR function by 1400 FDA-approved compounds in PDIO cultures of 76 different donors, covering 58 different CFTR genotypes. PDIOs from 47 donors were compound heterozygous for the F508del allele whilst PDIOs cultures from 29 donors harbored no F508del allele. Genotypes were stratified into different categories as has recently been published ^8^. Mutations not included in this study were large deletions or splicing mutations in close proximity of the splice site and were therefore categorized as Class I mutations, except for mutation A559T that was recently described to result in poor apical trafficking due to a defective folding of the CFTR protein and was therefore categorized as Class II ^11^. A list of all PDIOs and corresponding genotypes is provided in **Sup. Table 1**. PDIOs from 26 donors were compound heterozygous for a premature termination codon (PTC) mutation, PDIOs from 24 donors were compound heterozygous for a missense mutation, a splice mutation or a deletion and PDIOs from 5 donors had an insertion or an unclassified mutation **(Fig. 1A, left)**. All CFTR mutation classes are represented in our cohort except for Class III mutations and 6% of all alleles were unclassified **(Fig. 1A, right)**.

**Figure 1.**
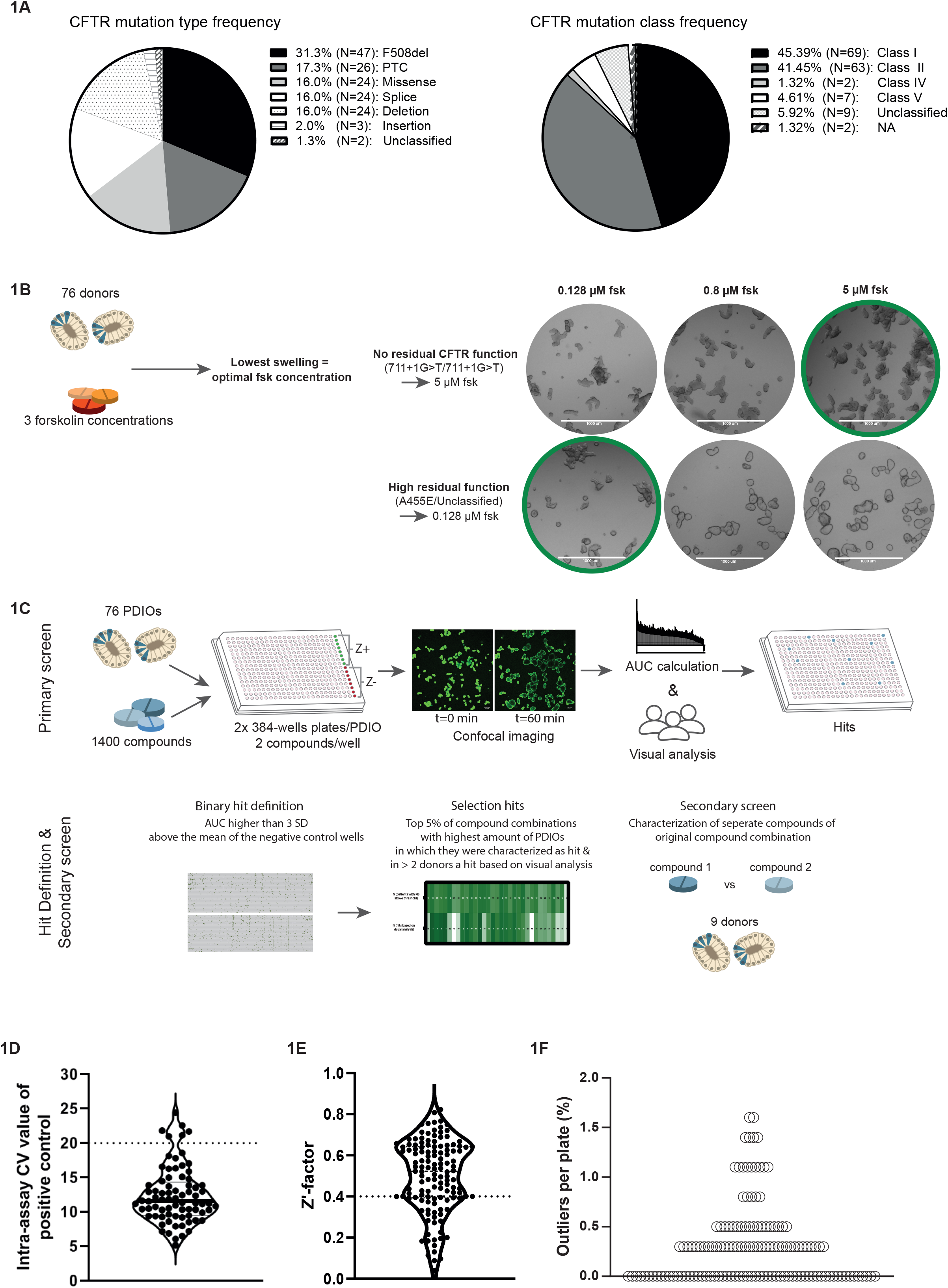
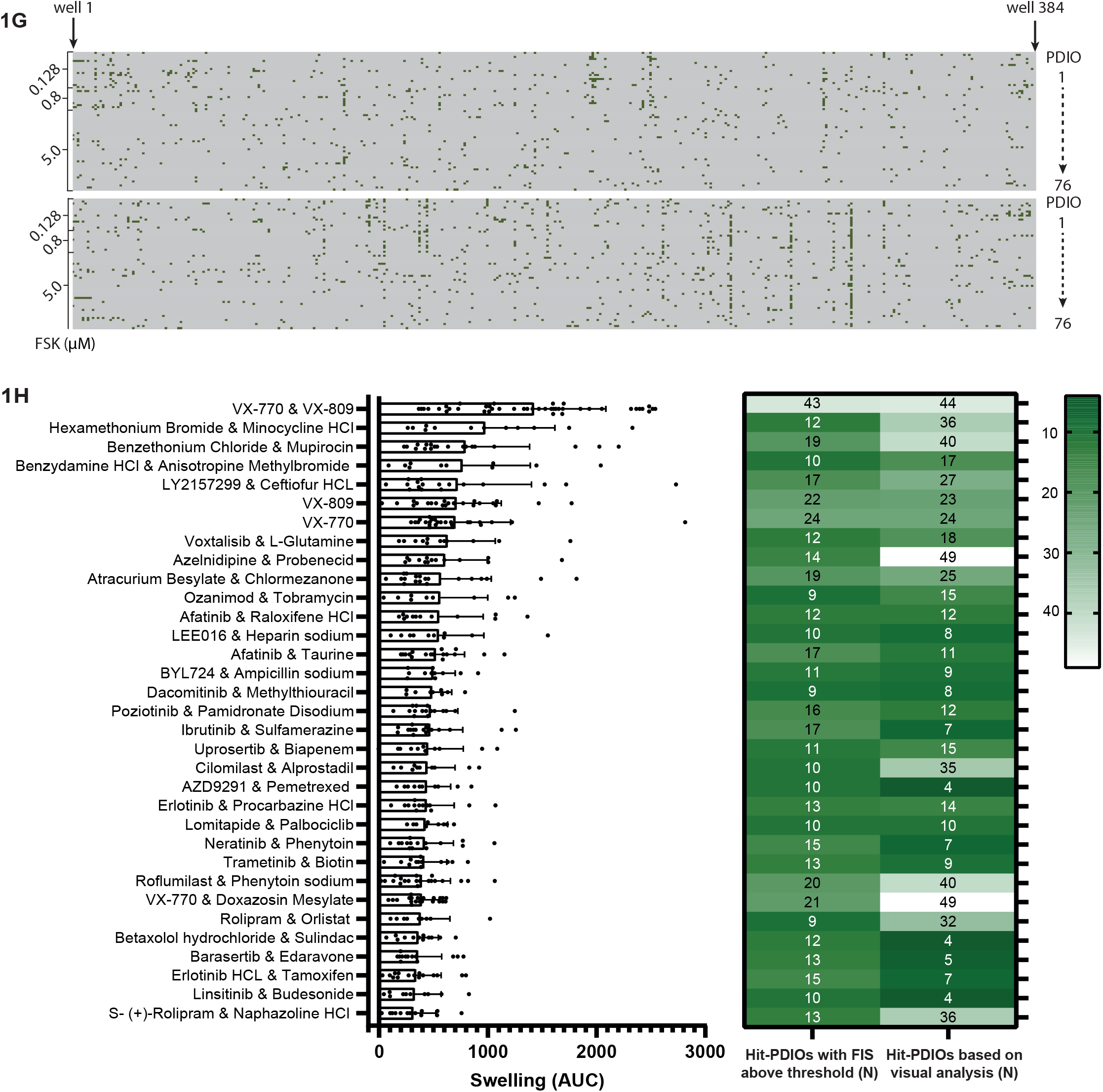
FDA-approved drugs increase FIS in non-homozygous F508del PDIOs. **(A)** Frequency of represented CFTR mutation types (left) and classes (right) of the 76 PDIOs included in the primary FDA screen. **(B)** Schematic of selection pipeline of forskolin concentration, based on residual CFTR function. Two PDIOs are shown as example that hold low (top) or low (boom) residual CFTR function resulting in respectively a high or low forskolin concentration during the FIS assays. **(C)** Schema-c of primary FDA screen describing PDIO plating, data analysis and decision pipeline for inclusion in secondary screen. **(D)** The intra-assay CV values of all plates, based on the Z+ values of all plates. The dotted line at 20 represent the upper limit value that indicates sufficient assay robustness. **(E)** The mean Z’-factor of all plates. The dotted line at 0.4 indicates a robust Z’-factor. **(F)** Outlier percentage of all plates. AUC values above >Q3+(3xIQR) (=5963) of all positive control wells or below Q1-(3xIQR) (=-452) of all negative control wells of the plates with a Z’-factor >0.5, were defined as outlier. **(G)** Binary outcome (hit or no hit) of all wells (from left to right) and PDIOs (from top to boom) divided over the two 384-wells plates. A well was selected as hit if the AUC was higher than the mean+3SD of the 8 negative control wells (DMSO treated) per 384-wells plate. Wells that were defined as hit are highlighted in green. **(H)** Overview of AUC values of the top 30 compound combinations and the three positive controls in primary screen, depicted for the PDIOs in which the compound combination was defined as hit. In the heatmap, numbers of PDIOs in which the compound combination was scored as hit based on AUC calculations are stated on the left, whereas numbers of PDIOs in which the compound combination was scored as hit based on visual analysis are stated on the right. Bars represent the means of all donors based on one technical replicate in one experiment per compound combination.

One challenge with screening a large variety of PDIOs is variation in baseline FIS due to differences in residual CFTR function. To compensate for this variability, we first determined the appropriate forskolin concentration per individual PDIO that resulted in the lowest level of baseline swelling, as to increase the chance of detecting compound-induced FIS per PDIO **(Fig. 1B)**. Based on visual analysis, we selected the forskolin concentration that resulted in the lowest amount of organoid swelling, resulting in 0.128 µM forskolin for PDIOs with high residual CFTR function (25% of all PDIOs), 0.8 µM forskolin in case of moderate residual CFTR function (15% of all PDIOs) and 5.0 µM forskolin in case of minimal or absent baseline swelling (60% of all PDIOs).

The screening pipeline consisted of a) PDIO addition to two 384-wells per PDIO, b) addition of two compounds per well for 24 hours, c) addition of forskolin directly before FIS measurements and d) confocal FIS measurements to visualize organoid swelling **(Fig. 1C)**. After FIS quantification and hit definition in a binary way (mean+3SD above negative control), we selected the compound combinations that were a) a hit in at least 12.5% (equal to 9 donors) of all donors based on AUC analysis and b) a hit in at least two donors based on visual confirmation. The individual compounds of this final list of hit compound combinations were evaluated in a secondary screen on 9 PDIOs representing various classes of CFTR mutations.

In the primary screen, F508del/S1252N organoids treated with 5 µM forskolin or with 5 µM forskolin + 3 µM VX-770 were used as positive control on each plate, negative controls were the PDIOs in question with forskolin. These conditions allowed verification of CV value calculation, Z’ factor calculation and outlier percentage calculation. CV values should not exceed 20% and CV values under 10% are considered excellent ^12^. The average CV value of 12.4% of all plates underlined assay robustness **(Fig. 1D)**. Additionally, Z’-factors were calculated as indicator of assay quality. The Z’-factor is a parameter based on positive and negative control that ranges between 0 and 1, with 1 indicating a perfect assay and Z’-factors larger than 0.4 considered acceptable ^12^. The average Z’-factor over all donors was 0.49, underlining the overall assay robustness **(Fig. 1E)**. 13 out of the 152 plates were excluded for hit selection. Eight were excluded due to image acquisition related technical errors, two because positive and negative controls were missing and three due to poor organoid quality. We removed outliers based on interquartile range (IQR) calculations where wells with AUC values above Q3+(3xIQR) (=5963) of all positive control wells or below Q1-(3xIQR) (=-452) of all negative control wells were excluded. All plates used for hit selection displayed an outlier percentage below 2% **(Fig. 1F)**.

Positive hits were selected based on FIS values that were higher than the mean+3SD of the 8 negative control wells within the identical plate **(Fig. 1G)**. The top 5% hits that increased AUC above this threshold in most patients, corresponded to 37 hits including the positive controls (VX-770, VX-809 and VX-770/VX-809). We selected the compound combinations that were a) a hit in at least 12.5% (equal to 9) of all donors based on AUC analysis and b) a hit in at least two donors based on visual analysis, resulting in a total of 30 compound combinations excluding positive controls, to be tested in the secondary screen. In **Fig. 1H**, the 30 top compound combinations and the three positive controls consisting of CFTR modulators are listed. For each compound combination the average AUC is stated, based on the AUCs of the PDIOs in which the compound combination was classified as hit, as well as the total number of PDIOs in which the compound combination was classified as hit and lastly the number of PDIOs in which a compound combination was distinguished as hit based on visual analysis. Overall, the three approaches of hit selection yield similar results. We observed large differences between PDIOS with respect to the amount of hits, where PDIO_047 with genotype F508del/G461R was responsive to a high number of compound combinations, in comparison to PDIO_069 with genotype R1162X/3539del16 for which no compound combinations were able to increase CFTR function **(Supplemental Table 2)**. Overall, the median number of hits differed per mutation category, ranging from 10.5 hits in Class II/Class V PDIOs to a median of 73 hits for Class II/Na or unclassified PDIOs **(Sup. Fig. 1)**. Additionally, the mean number of hits in the Class I/Class I category was significantly lower than the mean averaged number of hits of all categories combined (p=0.0467).

### Identification of Three Main Compound Families that Increase CFTR Function

We next set out to determine which of the two compounds of the selected wells was associated with the observed efficacy. PDIOs with 9 different genotypes were selected that represented the different CFTR mutation classes as well as the three different forskolin concentrations, representing different baseline CFTR function levels. PDIOs were treated with the individual 60 FDA compounds from the 30 original compound combinations. For most compound combinations, one of each compounds clearly resulted in a higher increase of FIS than the other compound. Z-scores were calculated in order to correct for differences between plates, compound with a Z-score higher than 1.5 in at least two donors were considered a hit, resulting in confirmation of 19 hits **(Fig. 2A)**. As anticipated, donor variation was observed between the selected hits, e.g. CFTR potentiator Ivacaftor on plate 1 (compound 2) reached a high Z-score across most genotypes, whereas Rolipram on plate 1 (compound 11) reached high Z-scores particularly in PDIOs with high residual function that were tested with 0.128 µM forskolin. Four compounds reached a Z-factor of 1.5 only in one donor, and in 9 compound combinations neither of both compounds was identified as hit. Average DMSO-corrected AUC levels of these 19 confirmed hits were calculated for the PDIOs in which the compound was defined as hit **(Fig. 2B)**. Interestingly, 3 main compound families could be identified. Firstly, CFTR modulator VX-770 resulted in CFTR rescue in genotypes for which this modulator is currently not approved. Additionally, 4 out of the 19 compounds were phosphodiesterase (PDE) inhibitors and 6 out of the 19 compounds were tyrosine kinase inhibitors (TKIs), that mainly inhibit EGFR. Due to the toxic nature of the latter category and the fact that PDE4 inhibitors are already used for airway disease COPD ^10,13^, we argued that PDE4 inhibitors and CFTR modulators were the most promising candidates.

**Figure 2.**
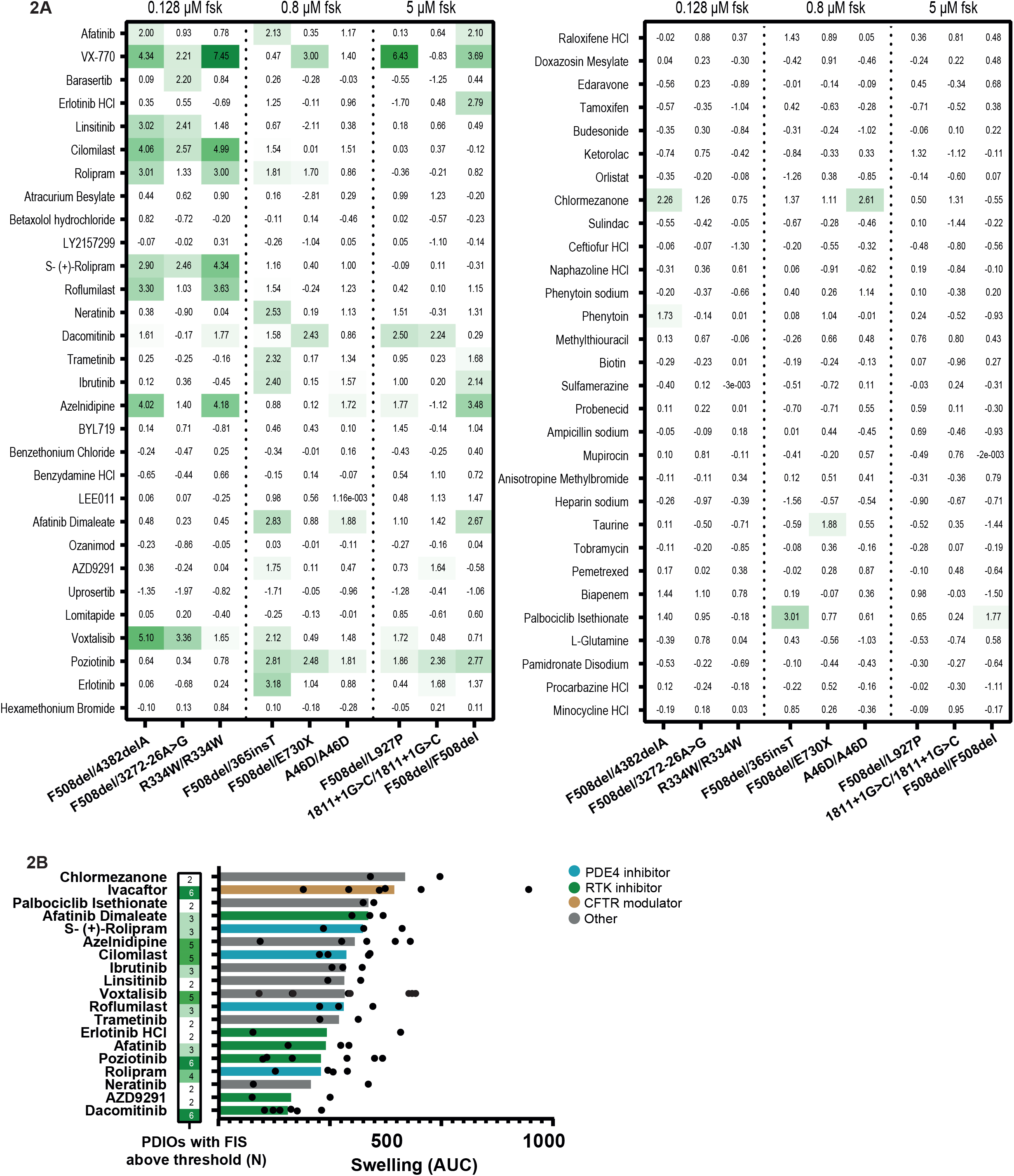
Idenfication of three main compound families that increase CFTR function. **(A)** Z-scores of secondary FIS screen of 9 PDIOs treated separately with the compounds of the top 30 compound combinations, based on one biological replicate experiment with one technical replicate. **(B)** The number of PDIOs in which FIS led to a Z-score above a threshold of 1.5 (left) and DMSO normalized AUC values of those PDIOs in which the compound was classified as a hit (right). Compound class is indicated by color, distinguishing between PDE4 inhibitors, RTK inhibitors, CFTR modulators and compounds with a distinct MoA.

### Acute PDE4 Inhibition Increases CFTR Function at a Low Nanomolar EC50

Phosphodiesterases (PDEs) comprise a group of enzymes that catalyze the hydrolysis of phosphodiester bonds of second messengers, cAMP and cyclic guanosine monophosphate (cGMP), thereby regulating many downstream signaling processes ^14^. PDE4 inhibitors act by blocking the catalytic site of PDE4, thereby suppressing cAMP degradation. Consequentially, intracellular cAMP levels rise thereby increasing PKA activation and subsequently increasing CFTR phosphorylation and function **(Fig. 3A)**. RNA expression analysis of the four PDE4 subtypes indicates identical expression between wild-type and CF (F508del/F508del) intestinal PDIOs. When comparing the PDE subtypes, higher expression of specifically PDE4D was observed in primary airway epithelial cells **(Fig. 3B)**.

**Figure 3.**
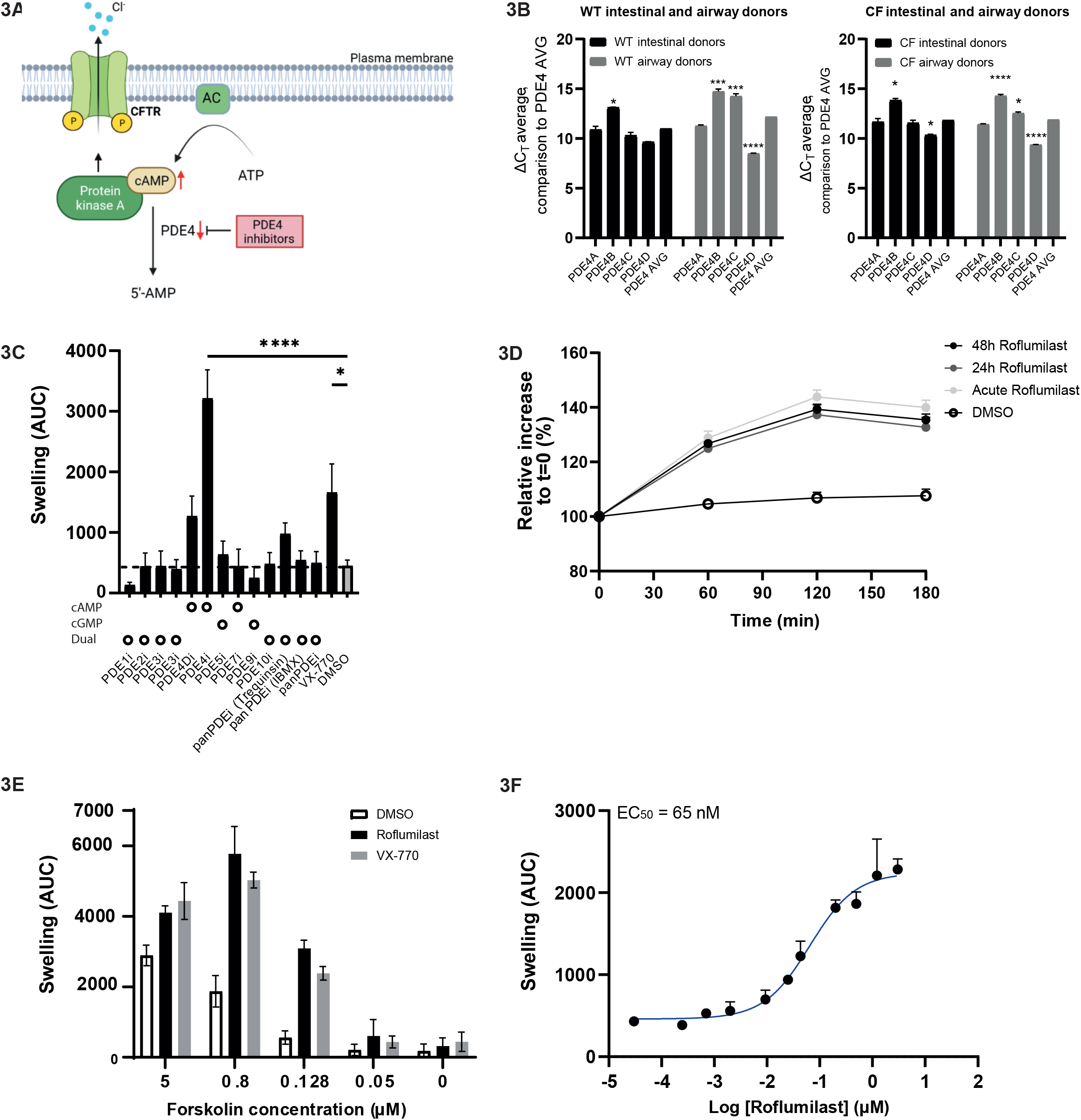
Acute PDE4 inhibition increases CFTR function at a low nanomolar EC50. **(A)** Schematic of mode of action of PDE4 inhibitors, that by suppressing cAMP degradation, result in subsequent elevation of intracellular cAMP, PKA activation and increase of CFTR phosphorylation and function. **(B)** Delta-CT values obtained via RT-qPCR of the diffierent PDE4 subtypes, normalized for CT values of housekeeping gene GAPDH, for WT and CF (F508del/F508del) donors in PDIOs and patient-derived, diffierentiated nasal epithelial cells. To calculate which PDE4 subtypes diffiered significantly from the average of all PDE subtypes, One-Way ANOVAs were performed for the CF/WT/intestinal/airway samples separately, followed by Dunnetts post-hoc analysis. Significance indicated by one asterisk corresponds to a p-value <0.05, significance indicated by three or four asterisks correspond to p<0.005 and p<0.0001, respectively. Bars indicate the mean of three technical replicates, derived of one biological replicate with errorbars indicating the SD. **(C)** FIS levels (AUC) of R334W/R334W PDIOs upon treatment with a range of PDE inhibitors. To assess significance, a One-Way ANOVA was performed followed by Dunnetts post-hoc analysis. Significance indicated by one asterisk corresponds to a p-value <0.05, significance indicated by four asterisks corresponds to a p-value <0.0001. Bars indicate the mean of three technical replicates, derived of three biological replicates with errorbars indicating the SEM. **(D)** Relative size increase during a FIS assay over me for R334W/R334W PDIOs upon preincubation or acute treatment with PDE4 inhibitor roflumilast (RF). Dots indicate the mean of three technical replicates, derived of three biological replicates with errorbars indicating the SEM. **(E)** FIS levels (AUC) of R334W/R334W PDIOs, upon acute treatment of RF and VX-770 and a titration range of forskolin. Bars indicate the mean of three technical replicates, derived of three biological replicates with errorbars indicating the SEM. **(F)** FIS levels (AUC) of R334W/R334W PDIOs, upon a concentration range of acute stimulation with RF. The EC50 was calculated based on logarithmic curve-fitting using GraphPad Prism and corresponds to 65 nM. Bars indicate the mean of three technical replicates, derived of three biological replicates with errorbars indicating the SEM.

We first studied selectivity of PDE subtypes for modulating FIS in R334W/R334W PDIOs, one of the PDIOs that was most responsive to PDE4 inhibitors based on the results of the secondary screen. Comparison of different PDE inhibitors that target cAMP or cGMP as well as three pan-PDE inhibitors, indicate that inhibition of mainly PDE4D results in a large increase of CFTR function **(Fig. 3C)**. Inhibition of other cAMP-mediated PDEs or cGMP-mediated PDEs did not increase of CFTR function. Pan-PDE inhibitors were less efficacious than PDE4-selective inhibitors. We continued with optimizing the dynamic range to detect effects of PDE4 inhibitors and observed that PDE4 inhibitor-induced swelling was at its maximum after two hours of incubation **(Fig. 3D)**. In the primary and secondary screen, all compounds were preincubated for 24 hours prior to FIS measurements. Acute addition and longer exposures of PDE4 inhibitors however all resulted in similar increases in FIS, consistent with the established mode-of-action of PDE4i as direct inhibitor of cAMP degradation **(Fig. 3D)**. This is distinct from β2 adrenergic receptors (β2AR)-agonists that increase cAMP levels and downregulate PDIO responses to forskolin. Especially upon longer exposure it has been shown that B2AR-agonist salbutamol pretreatment can result in reduced CFTR activity ^15^. We also detected a decrease in organoid swelling after 72h of pre-incubation with both salbutamol and roflumilast when compared with PDIO swelling induced by acute administration **(Sup. Fig. 2A)**. We however observed forskolin-independent PDIO preswelling prior to the FIS assay, indicated by an increase of steady-state lumen area (SLA) **(Sup. Fig. 2B)**. This potentially explains a decrease of PDIO swelling during the FIS assay. We subsequently calculated ratios between FIS decrease and SLA increase, where ratios larger than 1 indicate that the decrease of PDIO swelling after 72h incubation with salbutamol is larger than what is caused by PDIO preswelling. Whilst this is the case for salbutamol, this is not the case for especially the lower concentration of roflumilast **(Sup. Fig. 2C)**. This indicates that PDE4 inhibition, opposed to B2AR-agonist, does not result in downregulation of CFTR activity.

Without forskolin induced cAMP increase, PDE4 inhibition did not result in increased CFTR function. Additionally, when stimulating with a high concentration of forskolin, the difference between residual CFTR function and PDE4 induced CFTR function was not detectable due to high swelling in both conditions **(Fig. 3E)**. This underlines the forskolin dependency of the effect of PDE4 inhibitors as well as that 0.128 µM forskolin results in the optimal dynamic range to detect effects of PDE4 inhibition. Additionally, FIS was measured with increasing concentrations of roflumilast. A clear concentration-dependent effect was observed with an EC_50_ of 65 nM **(Fig. 3F)**.

### PDE4 Inhibition Efficacy Depends on Residual CFTR Function

To compare whether different PDE4 inhibitors result in differences in CFTR function increase, we compared 5 different PDE4 inhibitors: rolipram, roflumilast, cilomilast, piclamilast and apremilast. To further characterize genotype-specific effect, the PDE4 inhibitors were screened on a panel of 14 PDIOs. 8 PDIOs expressed different Class II/Class III mutations, 4 PDIOs were homozygous for the F508del/F508del CFTR mutation and 2 PDIOs homozygously expressed W1282X CFTR. Prior to characterizing CFTR function increase, we confirmed absence of toxicity of those PDE4 inhibitors **(Sup. Fig. 3)**. PDE4 inhibitors were tested alone or in combination with additional compounds. We compared compound-induced swelling to background-induced swelling, for which Pearson’s R^2^ correlations for roflumilast, rolipram and VX-770 are shown in **Fig. 4A**, and in **Sup. Fig. 4** for apremilast, cilomilast and piclamilast. Correlations were significant and positive for all compounds, with the highest for VX-770 (R^2^=0.95, p<0.0001) in comparison to roflumilast (R^2^=0.68, p<0.0001) and rolipram (R^2^=0.73, p<0.0001). Whilst this underlines that PDE4 inhibition efficacy is positively correlated to baseline CFTR function, we observed large variation in PDE4 inhibitor response between PDIOs with low residual function.

**Figure 4.**
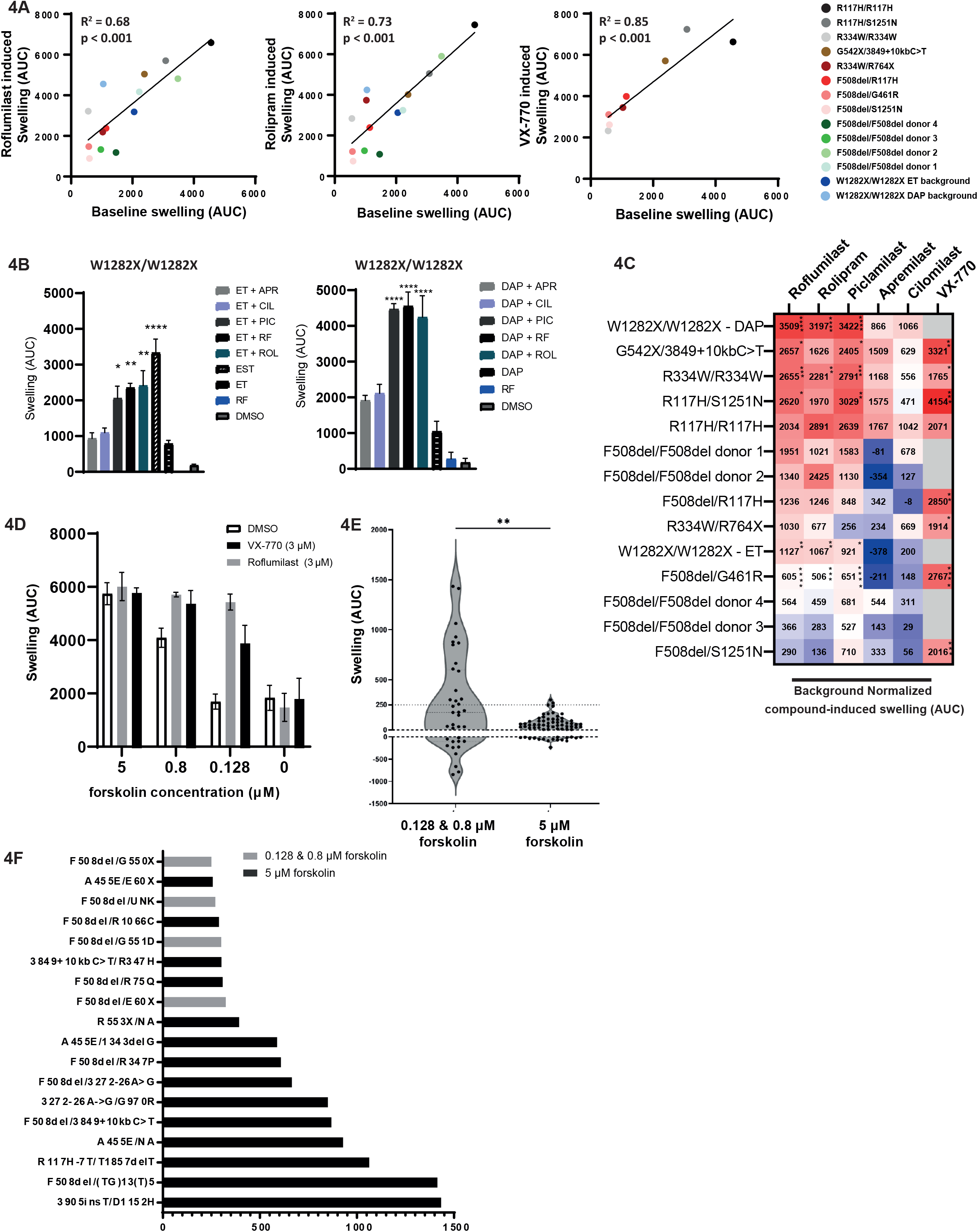
PDE4 inhibition efficacy depends on residual CFTR funticon. Roflumilast, rolipram or VX-770-induced swelling (AUC) versus background (DMSO/other) induced swelling (AUC), for 14 PDIOs indicated by the diffierent colored dots. Dots indicate the mean of three technical replicates, derived of three biological replicates. **(B)** W1282X-/W1282X PDIO swelling upon treatment with a panel of PDE4 inhibitors, in combination with ELX-02 (E), VX-445/VX-661/VX-770 (T) or SMG1i (S) (left) or DAP (right). Bars indicate the mean of three technical replicates, derived of three biological replicates with errorbars indicating the SEM. **(C)** A heatmap of compound-induced PDIO swelling, normalized to background compounds or DMSO. To calculate stascal significance, One-Way ANOVAs were performed per PDIO to compare compound-incuded swelling to baseline swelling, followed by Dunnes post-hoc tests. Values are the mean of three technical replicates, derived of three biological replicates and significant diffierences are depicted by one/two/three/four asterisks, corresponding to p-values smaller than 0.05, 0.01, 0.001 or 0.0001 respectively. **(D)** DMSO-corrected swelling of A445E/S1251N PDIOs upon treatment with roflumilast and a concentration range of forskolin. Bars indicate the mean of three technical replicates, derived of three biological replicates with errorbars indicating the SEM. **(E)** PDIO swelling of 107 PDIOs, treated with roflumilast, split into a group smulated with low forskolin (0.128 and 0.8 μM) or high forskolin concentrations (5 μM). To compare the groups treated with 0.128/0.8 μM forskolin to the groups treated with 5 μM forskolin, an unpaired two-tailed T-test was performed (p=0.0013). Dots indicate one replicate, derived of one biological replicates. **(F)** The 19 PDIOs in which, after DMSO normalizaon, an increase of >250 AUC was detected. Bars represent one replicate, derived of one biological replicate.

In total, in 6-out-of-8 non-Class I and non-F508del PDIOs, at least one PDE4 inhibitor significantly elevated CFTR function, in some cases resulting in AUC values similar or higher than VX-770 corrected positive control conditions **(Sup. Fig. 5A)**. PDIOs that respond better to PDE4 inhibitors than predicted based on the correlations, are the PDIOs harboring W1282X/W1282X CFTR. These PDIOs were treated with PDE4 inhibitors in combination with RT agent DAP **(Fig. 4B)**. The large additional effect of PDE4 inhibitors when combined with DAP, indicates compound synergy. This could be attributed to the MoA of DAP which results in tryptophan incorporation at the PTC site and therefore WT restoration of the CFTR protein ^16^. This is not the case for RT-agent ELX-02, for which we observed a less prominent increase in CFTR function when combined with PDE4 inhibitors & CFTR modulators Trikafta (VX-445/VX-661/VX-770). In this combination however, the combination of roflumilast/ELX-02/CFTR modulators reached PDIO swelling levels that were comparable to the combination of ELX-02/Trikafta/SMGi, the latter inhibiting nonsense-mediated mRNA decay (NMD). PDIOs that respond less to PDE4 inhibitors than predicted based on the correlation between background induced swelling and PDE4 induced swelling, are PDIOs homozygously expressing F508del/F508del CFTR that were tested in combination with CFTR modulators VX-809/VX-770 **(Sup. Fig. 5B)**. All responses are summarized in **Fig. 4C**, in which we show background-corrected AUC values upon PDE4 inhibitor treatment. The effect of PDE4 inhibition on PDIO swelling shows a large amount of variation, underlining genotype-associated differences in response. Overall, among all included PDIOs, a similar trend was observed regarding the effect of the different PDE4 inhibitors, with piclamilast, roflumilast and rolipram resulting in the highest increase in CFTR function.

**Figure 5.**
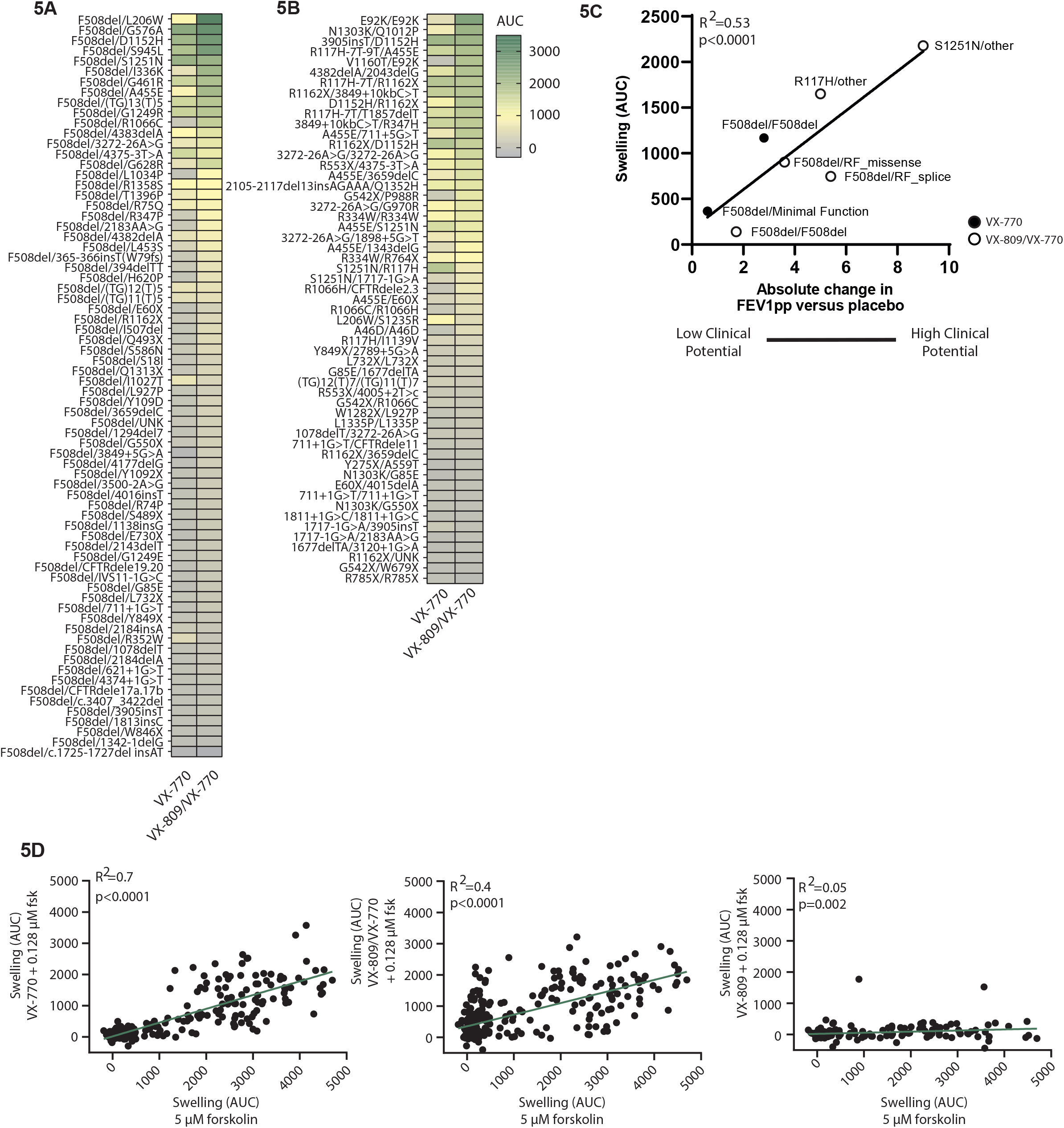
Potential of Label Expansion of CFTR modulators for People with Rare CFTR Genotypes. **(A-B)** VX-770 or VX-809/VX-770 induced swelling of PDIOs at 0.128 µM forskolin for **(A)** PDIOs expressing a F508del mutation on one allele and a rare CFTR mutations on the other allele and **(B)** PDIOs expressing two rare CFTR variants. Swelling is normalized for residual CFTR function by subtraction of DMSO-induced FIS. Values are based on one technical replicates, derived of one biological replicates. **(C)** Pearson correlation of drug-corrected PDIO swelling versus lung function increase (FEVpp). CFTR modulator swelling was measured at 0.128 µM forskolin and corrected for DMSO-induced swelling and is presented per CFTR genotype group. VX-770-treated PDIOs are represented by white dots whereas VX-809/VX-770 treated PDIOs are represented by black dots. FEV1pp versus placebo values are based on clinical trials, summarized in Table1. **(D)** Pearson correlation of PDIO swelling upon modulator therapy (VX-770, VX-809 or VX-809/VX770) and 0.128 µM forskolin versus 5 µM forskolin and DMSO-treated PDIOs.

We subsequently investigated the effect of PDE4 inhibition in primary airway organoids harboring A445E/S1251N CFTR, as these mutations have previously been recognized as Class II/III mutations respectively and possess some residual CFTR function. Corresponding to the results in the PDIOs, roflumilast elevates CFTR function in a forskolin dependent manner, where maximum efficacy is observed at mainly 0.128 µM forskolin **(Fig. 4D)**.

To stratify more CF patients that could potentially benefit from PDE4 inhibition, we tested roflumilast on 107 additional PDIOs, covering 74 genotypes of which 34 did not express F508del CFTR **(Supplemental Table 3)**. Roflumilast increased swelling (AUC>250) in 19 out of 107 PDIOs **(Fig. 4E)**. Genotypes that were responsive to treatment were confirmed, such as 3849+10kbC>T, and other responsive genotypes were identified, such as 3905insT/D1152H **(Fig. 4F)**.

### Potential of Label Expansion of CFTR modulators for People with Rare CFTR Genotypes

In the primary and secondary screen, CFTR modulators elevated CFTR function to a high degree and in a large number of PDIOs, suggesting a high potential for label extension of CFTR modulators. To further characterize genotypes that would potentially benefit from CFTR modulator therapy, we screened 197 PDIOs representing 127 genotypes, that carried at least one CFTR mutation that is present in <1% of the European and American population and carrying maximum one of the following alleles: F508del/G542X/G551D/R117H/N1303K/ W1282X/3849+10kbC>T/R553X/1717-1G>A/621+1G>T/2789+5G>A/3120+1G>A/CFTRdele2,3 **(Supplemental Table 4)**. Modulator responses of another 109 PDIOs **(Supplemental Table 5)** representing 34 different genotypes were additionally screened to obtain an overview of reference AUC levels allowing characterization of the correlation between FIS data and clinical data at group level. FIS was measured upon activation with 0.128 µM forskolin, as the *in vitro* drug effect expressed by FIS measured with this forskolin concentration has previously been shown to correlate with the *in vivo* drug effect.

The 197 PDIOs were treated with VX-770 or VX-809/VX-770. FIS data are shown for PDIOs carrying at least one F508del allele in **Fig. 5A**, and FIS data for non-F508del PDIOs are shown in **Fig. 5B**. As done before in a smaller dataset ^6^, we investigated the association between average FIS values in PDIOs and the average FEV1 response in clinical trials **(Table 1)** in 7 representative genotype-stratified subgroups **(Fig. 5C)**. Consistent with previous findings ^7^, we found a significant correlation (R^2^=0.53, p<0.0001) between the level of CFTR-modulator induced swelling of the PDIOs and the treatment effect expressed in absolute change in FEV1pp of reported clinical studies. These data indicate that 31 of the 127 genotypes had VX-770-responses beyond that of VX-770 treated F508del/splice PDIO and that 36 genotypes of the 127 genotypes had VX-770/VX-809 responses beyond that of VX-770/VX-809-treated F508del/F508del PDIO, indicating a clinical benefit based on the correlation described in **Fig. 5C**. Furthermore, PDIOs with CFTR mutations that are currently not categorized into one of the CFTR mutation classes, can be classified based on these data.

**Table 1:** Overview of clinical trials with CFTR-correcting treatments in subjects expressing different CFTR mutations. For the R117H trial, only data from CF subjects aged >18 were used, because subjects aged 6 to 18 had a different mean baseline FEV1 compared to those in the other trials. The numbers correlate with the numbers in **Fig. 5C**. NS: not significant, NA: statistical analysis not performed due to small numbers for individual mutations, RF: residual function, MF: minimal function.

| *Treatment* | *Genotype* | *Absolute change in FEV1pp versus placebo* |
| --- | --- | --- |
| 1. VX-770 | F508del/F508del | 1.72 (NS)<sup>37</sup> |
| 2. VX-770/VX-809 | F508del/MF | 0.6 (NS)<sup>38</sup> |
| 3. VX-770 | F508del/RF_splice | 5.4 (NA)<sup>39</sup> |
| 4. VX-770 | F508del/RF_missense | 3.6 (NA)<sup>39</sup> |
| 5. VX-770/VX-809 | F508del/F508del | 2.8 (p<0.001)<sup>40</sup> |
| 6. VX-770 | R117H/other | 5 (p=0.01)<sup>41</sup> |
| 7. VX-770 | S1251N/other | 9 (NA)<sup>42</sup> |

We observed a strong correlation (R^2^=0.7) between baseline swelling (DMSO) at 5 µM forskolin and swelling increase with VX-770 and 0.128 µM forskolin **(Fig. 5D**). A similar relation between residual CFTR function and VX-770/VX-809-mediated increase in swelling was observed, however with a lower R^2^ (R^2^=0.4). Only two organoid cultures (E92K/E92K and A455E/(TG)13(T)5) showed an increase in CFTR function with treatment of solely VX-809, and correlation between VX-809 induced swelling response and DMSO-induced swelling response was absent.

## DISCUSSION

Preclinical cell-based assays that recapitulate human disease can play an important role in the first steps of drug repurposing. We previously developed an *in vitro* functional assay using PDIOs to measure function of the CFTR gene that is mutated in pwCF. We screened 76 non-homozygous F508del-CFTR PDIOs to measure the efficacy of 1400 FDA-approved drugs on improving CFTR function as measured by FIS. The utilized FIS assay allows read-out of functional CFTR, which has been shown to associate with disease severity indicators of CF, long-term disease progression and therapeutic response, underlining the potential clinical value of identified preclinical hits ^7,8^. Here, we show that PDE4 inhibitors piclamilast, roflumilast and rolipram, are potent CFTR inducers in PDIOs where residual CFTR is either present, or created by additional compound exposure. Additionally, upon CFTR modulator treatment (VX-809/VX-770), we show rescue of PDIOs that are currently not eligible for this therapy.

The procedure to biobank and culture PDIOs for screening was robust. Previous optimization of several steps in 96-wells FIS assay ^4^, allowed us to practically perform the FIS assay in a 384-wells format ^9^ .In the context of high-throughput screenings, most efforts have been made in the cancer-field. Whilst patient-derived organoid screening has received much attention, many studies in this context are lower-throughput screenings, in which read-outs are often centered around viability ^17–19^. Whilst viability can be quantified in a relative straightforward fashion, for example by luminescence measurement, increasing throughput of functional assays with a more complicated read-out is exceedingly challenging. A few recent studies reported higher throughput screenings on 2D patient-derived material in the context of CF ^20,21^, yet robustness of exploited assays was either lower than in our study or not reported. Overall, robustness of our screening assay was confirmed and underlined by 70% of the plates reaching Z’-factors of 0.4 or higher and the average of all plates reaching a Z’-factor of 0.5. As a positive control was not available for all PDIOs, the Z’-factor positive signal on each plate consisted of F508del/S1251N PDIOs stimulated with VX-770 and forskolin. For this reason, we did not use Z’ calculations to exclude individual plates from the analysis. Further improvement of assay robustness and throughput might come from improving automation by means of automated organoid dispensers, drug printers and centrifugal washers to further reduce technical variability.

An additional challenge we encountered, is that PDIOs differ in baseline residual CFTR function, thereby limiting the opportunity to detect positive hits for individual PDIOs with high forskolin (optimal for low baseline CFTR) and low forskolin (optimal for high baseline CFTR) stimulation respectively. PDIO-specific forskolin concentrations were thus selected and enabled screening by minimizing baseline swelling, but prevented the use of a uniform assay for all PDIOs. Wild-type, non-CF organoids have pre-swollen phenotypes and show lower responses to forskolin due to their already fluid-filled lumens. Using FIS as screening readout, might have hampered detection of highly promising hits that caused significant PDIO pre-swelling prior to forskolin stimulation. We therefore visually inspected all wells of the stimulated PDIOs, but found no strong pre-swollen organoid phenotypes, apart from wells containing cAMP-increasing drugs like β2-agonists that we previously reported ^22^. In the future, new assays that are based on image analysis of absolute steady-state phenotypes need to be developed to complement the current kinetic assay that rely on relative changes in organoid phenotypes.

PDIO swelling is highly CFTR dependent under standard culturing conditions, and as such we anticipated that positive hits might either increase the CFTR apical protein expression, channel open probability or channel conductivity. The primary screen resulted in a list of 30 top compound combinations. Large differences between PDIOs were observed, where some were non-responsive overall and some were sensitive to a high number of compound combinations. Overall, PDIOs with two Class I mutations were the least responsive. To limit workload, we chose to validate the hit compound combinations in 9 PDIOs that represented the different CFTR mutation classes and baseline CFTR levels, as stratified by the three different forskolin concentrations. A limitation of this approach is that we did not fully recapitulate the patient and mutation variation of the initial screen, which potentially resulted in a loss of hit compounds in this validation screen. In the secondary screen, we showed that 19 compounds out of the 30 compound combinations resulted in an increase in FIS. Three main families were distinguished within these compounds, existing CFTR modulators; PDE4 inhibitors and tyrosine kinase inhibitors (TKIs). Due to the toxic nature of the latter category ^23,24^ the fact that PDE4 inhibitors are already used for smooth muscle relaxation in the respiratory disease COPD ^10^ and that for CFTR modulators it would be a matter of label extension instead of drug repurposing, we further investigated those latter two subfamilies further in our studies.

PDEs catalyze the hydrolysis of phosphodiester bonds of second messengers, cAMP and cyclic guanosine monophosphate (cGMP), thereby regulating many downstream signaling processes such as smooth muscle activation and inflammation associated pathways ^14^. By inhibiting smooth muscle activation, roflumilast is the first PDE4 inhibitor that has received regulatory approval for the treatment of a subset of patients with severe chronic obstructive pulmonary disease (COPD) ^25^. We verified that PDE4 is indeed the main PDE variant whose inhibition is related to CFTR function elevation, and found also higher expression of this PDE variant than of the other PDE variants in both PDIOs as well as primary nasal epithelial cells differentiated at air-liquid interface. Whilst PDE4 inhibition can increase CFTR activation due to higher levels of cAMP and subsequent PKA activation and increased CFTR channel opening, PDE4 inhibition does not restore CFTR function directly. This is underlined by the absence of PDE4i-mediated CFTR increase in W1282X/W1282X PDIOs when no other compounds are combined with the PDE4 inhibitors. Additionally, we show positive correlations between residual CFTR function and response to PDE4 inhibitors. Among all included PDIOs, piclamilast, roflumilast and rolipram elevated CFTR function to the highest extent. The difference between those PDE4 inhibitors and apremilast and cilomilast could be related to differences in the potency as well as PDE4 subtype selectivity. Roflumilast and piclamilast have previously been characterized by high subnanomolar potency with IC_50_ at 0.2-4.3 and 1 nM, respectively ^10,26^. On the other hand, IC_50_ of apremilast and cilomilast were identified at higher concentrations of 74 and 110 nM, respectively ^27,28^.

It was interesting to observe that there were large differences between the PDIOs and the extent of response to PDE4 inhibitors. Of the different PDIOs characterized in this study, several genotypes benefited from PDE4 inhibition as single compound, such as R334W, 3849+10kbC>T and G461R. We additionally assessed the effect of PDE4 inhibition in combination with additional compounds. We show that large synergistic effects can be achieved by combination of PDE4 inhibitors and compounds with different MOAs, such as DAP and roflumilast in W1282X/W1282X PDIOs. Strikingly, PDE4 inhibition did not further increase CFTR function in F508del/F508del PDIOs with either VX-809/VX-770 or other CFTR modulators (*data not shown*). Differences in the intracellular pathways such as low cAMP levels or differences in phosphorylation susceptibility caused by different compound treatments, might explain this absence of the PDE4 inhibitor-related effects. Characterizing cAMP and PKA levels or the degree of phosphorylation of CFTR in future studies could be of added value to further understand the difference between the F508del/F508del and the other genotypes. Despite the promising results of PDE4 inhibition in regards to elevation of CFTR function, we note that the *in vivo* efficacy of cAMP modulating pathways could be overestimated *in vitro* due to the differences in physiological cAMP concentrations *in vivo* and *in vitro*. Additionally, levels of baseline cAMP level potentially vary across tissues resulting in different, tissue-specific PDE4 effects.

Among all FDA hits, CFTR function modulators were most effective. Importantly, as CFTR modulators are already approved for specific mutations causing CF, it would be a matter of label extension instead of drug repurposing, which could result in an even faster translation into the clinic. Consistent with a previous study investigating this correlation ^6^, we found a significant correlation between the level of the DMSO-corrected drug-induced swelling of the PDIOs with 0.128 µM forskolin and the treatment effect expressed in absolute change in FEV1pp of available clinical trial data of mutations present in our study. Recently, the triple combination of CFTR modulators VX-445/VX-661/VX-809 has been approved by the FDA and EMA for all non-homozygous F508del genotypes. Concerning our screen on non-F508del PDIOs that are currently not approved for any modulators, we show based on our association between FIS data and clinical data that 17 out of 54 or 23 out of 54 PDIOs included in this dataset could have a moderate clinical benefit of respectively VX-770 or VX-809/VX-770 therapy, as their swelling response is equal or higher than the F508del/RF_Splice PDIO category treated with VX-770. The mutations 4382delA/2043delG and R334W/R334W are particularly interesting as these mutations are currently not approved for VX-770 therapy. These results underline the relevance of continuing to screen non-eligible non-F508del-CFTR genotypes with CFTR modulators and to potentially expand the label of these compounds based on the FIS assay. Additionally, our results and screening pipeline overall can aid in theratyping CFTR mutations of unknown consequence into a mutation category. For example, CF0823 (G542X/P988R) responds well to the combination of VX-770/VX-809 whilst swelling is not increased upon VX-770 treatment alone, indicating that mutation P988R is a CFTR mutation that results in improper CFTR folding and trafficking.

Additional to repurposing of PDE4 inhibitors and CFTR modulators, we found several other hit families that may reveal new targets and pathways acting on CFTR and that could be further characterized in the future. In the secondary screen, several TKIs were found to elevate CFTR function, such as Afatinib and Erlotinib. Interactions between CFTR and TKIs such as Afatinib have indeed previously been described, for example in the context of RTK inhibitor induced diarrhea ^29^. A recent study describes that EGFR TKIs potentiated the activity of potassium and CFTR chloride channels in T84 cell monolayers and rat models ^30^. However, as TKIs are mainly used as anti-cancer therapeutics and are known for severe side-effects, rapid translation of these results to the clinic is challenging. Voxtalisib, a PI3kinase and mTOR inhibitor additionally increased organoid swelling. Inhibitors of the PI3K/Akt/mTOR pathway have previously been shown to improve F508del-CFTR stability and function by stimulating autophagy in CFBE cells ^31^. Whether Voxtalisib acts with a similar MoA remains unclear for now. Another potentially interesting target we identified are GABA-activated chloride channels. Potentiation of the effects of the inhibitory neurotransmitter GABA with chlormezanone, also a compound among our hits, stimulates chloride influx through GABA-activated chloride channels ^32^. Although it is believed that the GABA receptor is predominantly expressed in the nervous system, some studies describe expression in intestinal epithelial cells and furthermore show involvement in intestinal fluid secretion ^33–35^. Potentially, future studies could aid in further characterization of the MoA of TKI/mTOR inhibition and GABA-inhibition mediated CFTR function increase and give leads for further drug development/biomedical chemistry based approaches.

In conclusion, we implemented a high-throughput 384-wells version of the functional FIS assay to screen a large number of PDIOs for compounds that enhance CFTR function. We characterized PDE4 inhibitors as novel CFTR elevating compound family, and furthermore show that CFTR modulators such as VX-809 and VX-770 might be beneficial for CF patients with CFTR mutations that are not eligible for CFTR modulators at present-day. We propose to conduct clinical studies designed to test the effects of roflumilast and existing CFTR modulators for these patients. Overall, our study demonstrates how preclinical studies using PDIOs can be used to initiate drug repurposing efforts. It facilitates the identification of potential treatments and responsive patients, thereby paving the way for patient stratification in the upcoming era of personalized medicine.

## MATERIALS AND METHODS

### Collection of primary epithelial cells of CF patients (pwCF)

All experimentation using human tissues described herein was approved by the medical ethical committee at University Medical Center Utrecht (UMCU; TcBio#14-008 and TcBio#16-586). Informed consent for tissue collection, generation, storage, and use of the organoids was obtained from all participating patients. Biobanked organoids are stored and catalogued (https://huborganoids.nl/) at the foundation Hubrecht Organoid Technology (http://hub4organoids.eu) and can be requested at

### Human intestinal organoid culture and forskolin selection

Patient-derived intestinal organoid (PDIO) culturing was executed as previously described ^4^. Prior to the FIS-assay, residual function levels of CFTR were determined during culture by visual analysis. Each PDIO culture was incubated with 0.02, 0.128, 0.8 and 5.0 µM forskolin for 1h, after which PDIO swelling was checked visually with a light-microscope. The forskolin concentration that resulted in lowest levels of residual swelling was chosen for subsequent screenings.

### Compounds

The FDA library, purchased from SelleckChem (Z178323-100uL-L1300), was stored at -80°C. All other compounds used in this study are listed in **Table 2**.

**Table 2:**
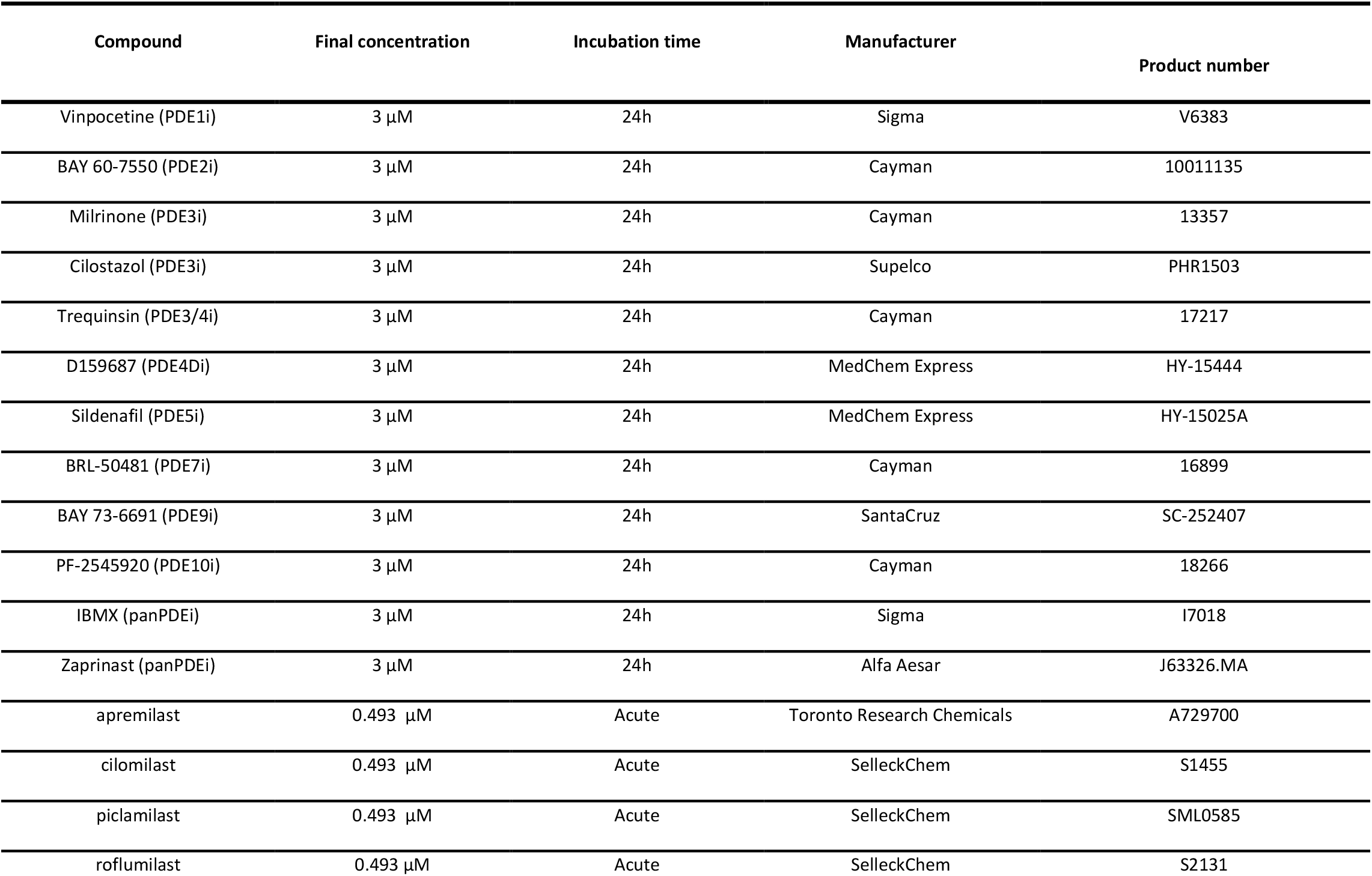
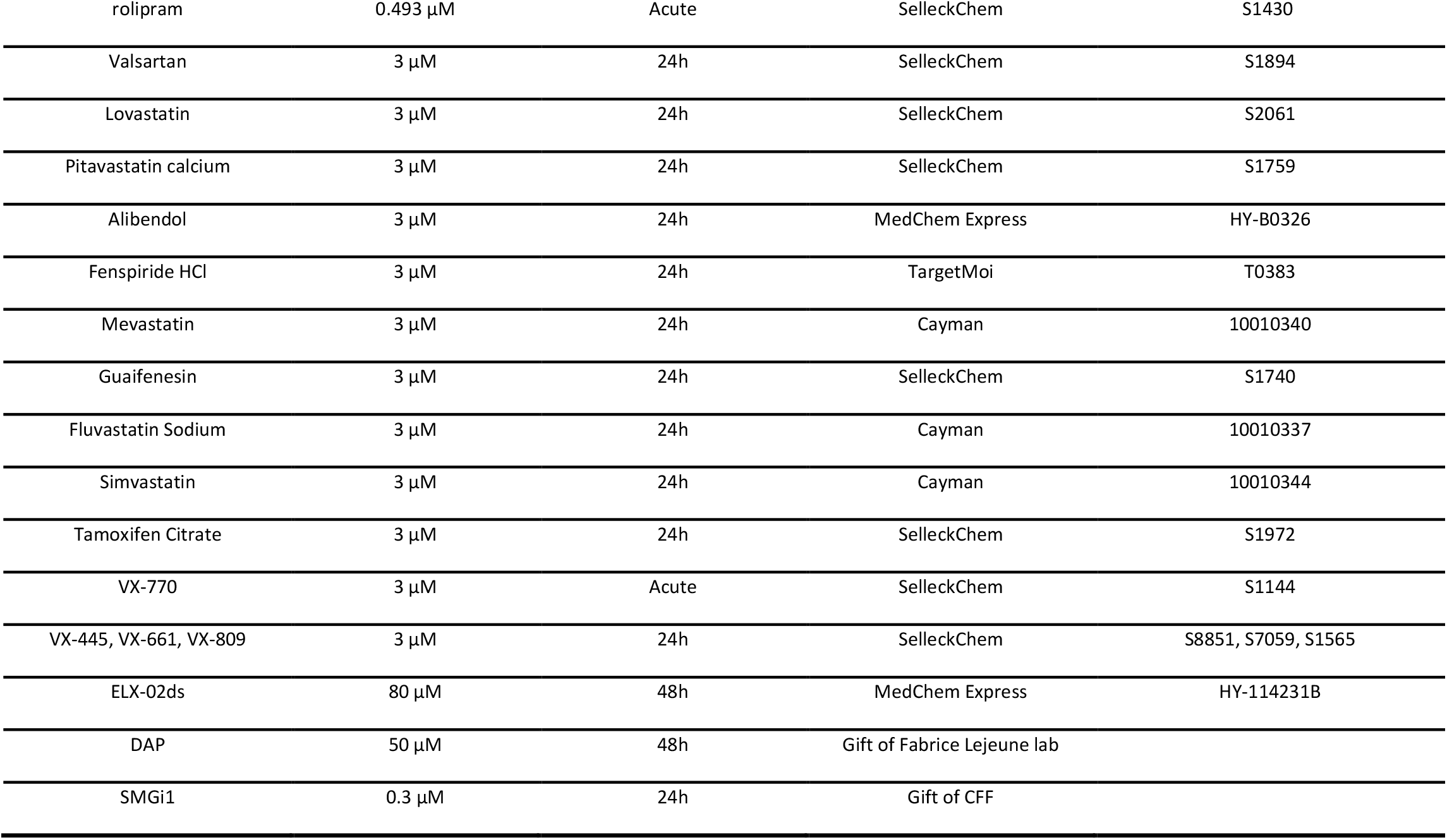
List of compounds used in this study

### 384-wells FIS assay

384-wells FIS-assays were performed according to previously described protocols ^4,5^, with minor adaptations allowing a 384-wells screening setting as summarized in **Table 3** ^9^. PDIOs of 76 different donors were seeded in 25% matrigel on two 384-wells plates/donor (7 µL/well). Organoids were subsequently submerged in 8 µL complete culture media supplemented with two FDA compounds/well (3 µM). The bottom 8 wells of the last column of each plate were not supplemented with FDA-compounds and served as negative control as well as minimal signal for Z’-factor calculations. The top 8 wells of the last column of each plate contained F508del/S1251N organoids that were treated with VX-770 (3 µM, acute addition) and forskolin (5 µM, acute addition), serving as a positive control and maximal signal for Z’-factor calculations. After 24 hours, 30 minutes prior to confocal imaging, organoids were fluorescently labeled with 5 µL calcein green (7 µM). 50 µL DMEM-F12 supplemented with forskolin and VX-770 for the positive controls, was added. Organoid swelling was monitored during 1 hour and total organoid surface area per well was quantified. Additional to fluorescent confocal images, brightfield images were taken of each well for visual analysis of organoid swelling. AUC values above 5963 (=Q3+(3xIQR) of all positive control wells of all plates with a Z’-factor > 5) and AUC values below -452 (=Q1-(3xIQR) of all negative control wells were excluded. Only plates with an outlier percentage below 2% were included for hit selection. Wells were selected as hit when AUC values were higher than the mean+3xSD of the 8 negative control wells (DMSO treated) on each individual plate. The top 5% hits that increased AUC above the threshold in most patients and that were a hit in at least 2 PDIOs based on visual analysis resulted in 33 compound combinations. The total number of hits we investigated in a secondary screen was 30 as three of the identified hits were the positive controls (VX-770, VX-809 and VX-770/VX-809). Since in the primary screen two compounds per well were combined, the secondary screen consisted of in total 60 individual compounds. The primary and secondary screen were performed once with one technical replicate per condition, except for negative and positive controls (8 replicates each).

**Table 3:** List of adaptations from the original 96-wells FIS assay protocol to allow 384-wells FIS screening

|  | 96-wells format, as described by <sup>4</sup> | 384-wells format, as described by <sup>9</sup> |
| --- | --- | --- |
| 1 | Plates are prewarmed at 37°C prior to organoid-matrigel addition | Plates are precooled at -20°C, and kept on ice during organoid-matrigel plating |
| 2 | 4 μl 50% matrigel organoid suspension is added per well as drops | 10 μl 25% matrigel organoid suspension is added per well with an automatic multichannel, to cover the whole surface |
| 3 | - | Plate is spun down in a centrifuge to ensure matrigel coverage of the whole well surface |
| 4 | After 10 minutes at 37°C, 50 μl medium per well is added | After 10 minutes at 37°C, 8 μl medium per well is added |
| 5 | X/Y/Z location is manually set for each well prior to image acquisition | X/Y/Z is automatically set based on plate lay-out and autofocus of anchor points during image acquisition |

### 96-wells FIS assay

96-wells FIS assays were conducted as previously described ^4^. For the secondary FDA screen, PDIOs derived from 9 donors were seeded into 96-wells plates within 50% matrigel. PDIOs were submerged in complete culture medium supplemented with one of the FDA compounds (3 µM), except for three wells (only DMSO). Z-scores of the secondary screen were determined according to the following formula: z-score = (x-μ)/σ, where x is the AUC value of each condition, μ is the mean AUC value of the 3 control wells on each plate, and σ is the standard deviation of the same 3 control wells on each plate. Besides the negative controls, each plate contained two positive control wells with F508del/S1251N organoids that were treated with VX-770 and 5 µM forskolin. For each donor, a suboptimal forskolin concentration was used, i.e. a forskolin concentration that resulted in minimal PDIO swelling. Organoid swelling was monitored during 1 hour and total organoid surface area per well was quantified ^7^.

For all follow-up FIS experiments on PDE4 inhibitors after the primary/secondary screen, PDE4 inhibitors were added acutely prior to the FIS measurement in combination with 0.128 µM forskolin prior to a 2hr measurement FIS assay, except when stated otherwise. Three biological replicate experiments were performed with three technical replicates per condition.

The screening of 107 different organoid cultures upon roflumilast treatment was assessed with 24h of roflumilast preincubation, and a donor-dependent suboptimal forskolin concentration (either 0.128, 0.8 or 5.0 µM) was used for the FIS-assay. This screen was performed once with one technical replicate per condition.

For the CFTR modulator screen, CFTR modulators VX-770 (3 µM, simultaneously added with forskolin), VX-809 (3 µM, 24h) and VX-770/VX-809 were tested on an additional 236 cultures, covering 167 different genotypes **(Supplemental Table 4)**. Prior to the 1h FIS measurements, CFTR activation was stimulated by addition of 0.128 µM forskolin for all genotypes. Screening was performed once with one technical replicate per condition.

### PDIO viability

Cell viability was assessed by means of an Alamar Blue assay performed on the PDIOs in the FIS assay plate, after the FIS assay ended. Organoids were treated with the PDE4 inhibitors or salbutamol at the indicated concentrations and incubation times. PDIOs were incubated with Alamar Blue (1:10 diluted in DMEM/F12 phenol-red free) for 4h at 37°C. Fluorescence intensity of the Alamar Blue solution was measured with a photo spectrometer at 544/570 nm. Viability was normalized to the averages of the positive (10% DMSO) and negative controls. Three biological replicate experiments were performed with three technical replicates per condition.

### PDIO lumen size

Confocal images obtained in the FIS assay results were used for the quantification of organoid lumen area and subsequently drug-induced swelling prior to the FIS assay. The luminal area as well as the total area was quantified manually using ImageJ, in a blinded fashion by 2 researchers. Results from three wells were averaged prior to calculation of the percentage of luminal organoid surface area of the total organoid surface area. Three biological replicate experiments were performed in which 10 organoid structures were characterized per condition.

### PDE4 quantative RT-qPCR

Prior to qPCR, total RNA was isolated from the airway and intestinal organoids using 350 µl RNeasy lysis buffer. RNA extraction was performed using the RNeasy Kit according to the manufacturer’s instructions and RNA yield was determined by a Nanodrop spectrophotometer. Subsequently, cDNA was synthesized using an iScript cDNA synthesis kit according to the manufacturer’s protocol. Next, 10 μl qRT-PCR reactions were executed using BIO-RAD I-Cycler 96 wells-plates with iQ™ SYBR Green Supermix and 10 μM forward and reverse primers. The samples were incubated for 3 minutes at 95 °C and 39 cycles at 10 seconds at 95 °C and 30 seconds at 62 °C. For the expression levels of PDE4 enzymes, ΔCt values were calculated while for the treated PTC organoids ΔΔCt values were calculated. The Ct values were normalized with the mean of mRNA expression of YWHAZ and GAPDH that served as housekeeping genes. Averages were calculated from the three technical replicates corresponding to one biological replicate. Melting peaks were analyzed to confirm specific primer binding. Details of primers used for qPCR are listed in **Table 4**.

**Table 4:**
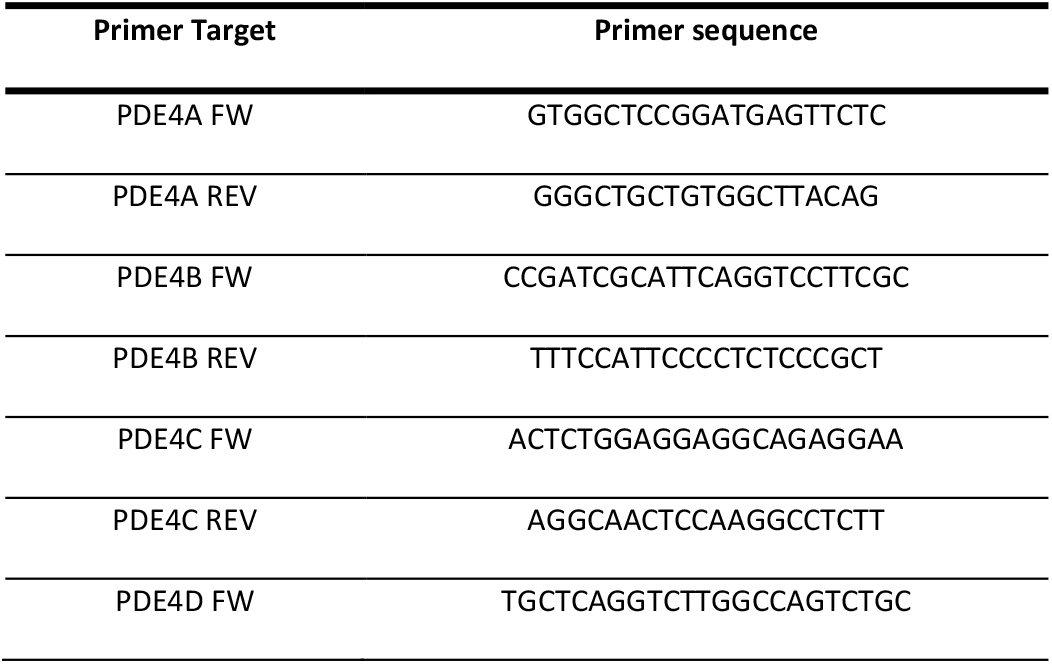
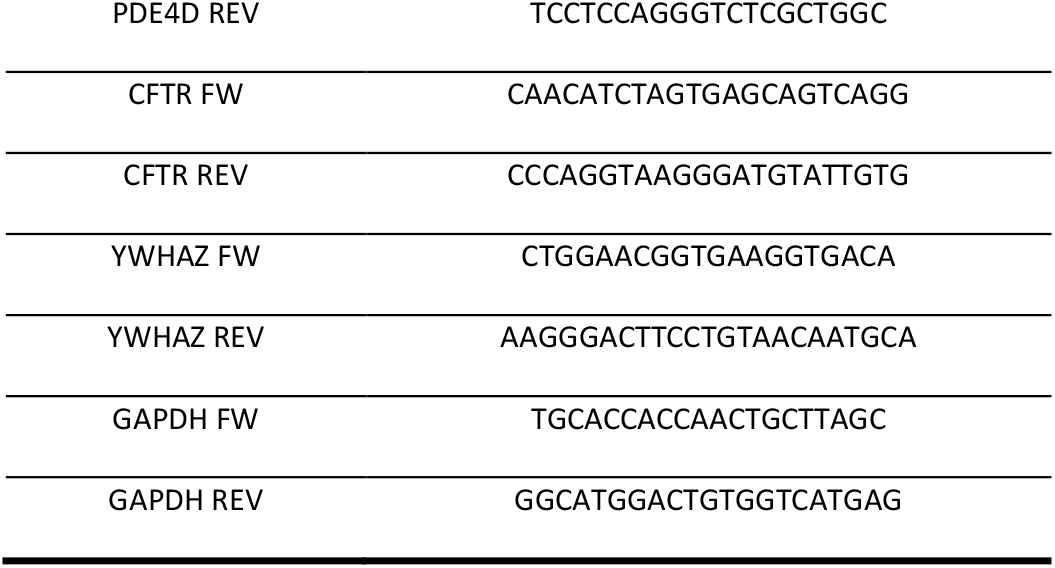
List of primers used in this study.

### Primary airway organoid FIS

FIS of primary airway organoids was performed as previously described ^36^. In brief, human nasal epithelial cells of early passage were cultured on 12-transwell inserts, previously coated with PureCol (1:100; 30 μg/ml) in expansion medium. When confluency was reached, culture medium was changed to air-liquid interface (ALI) differentiation medium supplemented with A83-01 in submerged condition for 2-3 days. Next, cells were air-exposed and further differentiated as ALI-cultures, refreshed at the basolateral side with ALI-diff medium supplemented with A83-01 and neuregulin-1β (NR, 0.5 nM). After 2-4 days, cells were refreshed with ALI-diff medium only with NR and without additional A83-01 and were differentiated for 3 weeks. The apical side of the cultures was washed with PBS once per week while the medium was refreshed twice a week. Upon 3 weeks of differentiation, organoid swelling was assessed in FIS assays, similar to the 96-wells PDIO FIS assays as described above. Averages were calculated from three technical replicates derived of three biological replicates.

### Statistical analysis

Statistical analyses were performed using GraphPad Prism®. For analysis of qPCR, One-Way ANOVAs were performed for the CF/WT/intestinal/airway groups separately to compare PDE4-subtype expression to the average expression of all PDE4 subtypes, followed by Dunnetts post-hoc analysis. For analysis of the PDE screen, a One-Way ANOVA was performed to compare swelling to DMSO control, followed by Dunnetts post-hoc analysis. Unless stated otherwise, graphs represent the average of 3 biological replicates which are obtained by averaging 3 technical replicates. To calculate statistical significance in the PDE4-screen, One-Way ANOVAs were performed per PDIO to compare compound-incuded swelling to baseline swelling, followed by Dunnetts post-hoc tests. When comparing two groups to each other, unpaired two-tailed T-tests were performed.

## DATA AVAILABILITY

Upon publication, data is available upon request via an online repository (DataverseNL).

## ACKNOWLEDGEMENTS

We would like to thank the people with CF who gave informed consent for generating and testing their individual organoids; all members of the research teams of the Dutch CF clinics that contributed to this work; and all colleagues of the HUB Organoid Technology for their help with generating intestinal organoid lines.

## FINANCIAL SUPPORT

This work was funded by grants of the Dutch Cystic Fibrosis Foundation (NCFS) as part of the HIT-CF Program and by ZonMW grant number: 91214103.

## AUTHOR CONTRIBUTION STATEMENT

E.d.P. and S.S. contributed to the design of the study, the acquisition, verification, analysis and interpretation of the data and have drafted the manuscript. P.V.M., G.N.I., S.W.F.S., A.M.V., J.E.B., E.K., H.O., M.C.H., G.B., K.M.d.W-d.G., S.H.-M., S.R.J., H.v.P., M.M.v.d.E., R.v.d.M., J.R., E.D., E.J.M.W., A.R.B., J.M.K. and G.H.K. contributed to the acquisition of study data and revised the manuscript. C.K.v.d.E and J.M.B. have made substantial contributions to the conception and design of the study, interpretation of data and revised the manuscript.

## DECLARATION OF INTEREST

J.M.B. reports personal fees from Vertex Pharmaceuticals, Proteostasis Therapeutics, Eloxx Pharmaceuticals, Teva Pharmaceutical Industries and Galapagos, outside the submitted work; In addition, J.M.B. has a patent patent(s) related to the FIS-assay with royalties paid. C.K.v.d.E. reports grants from GSK, grants from Nutricia, TEVA, Gilead, Vertex, ProQR, Proteostasis, Galapagos NV and Eloxx outside the submitted work; In addition, C.K.v.d.E. has a patent 10006904 with royalties paid. G.H.K. reports grants from Lung Foundation of the Netherlands, Vertex Pharmaceuticals, UBBO EMMIUS foundation, GSK, TEVA the Netherlands, ZON-MW (Vicigrant), European Union (H2020), outside the submitted work; and he has participated in advisory boards meetings to GSK and PURE-IMS outside the submitted work (Money to institution). All other authors have nothing to disclose.

---

**Sup. Fig. 1.**
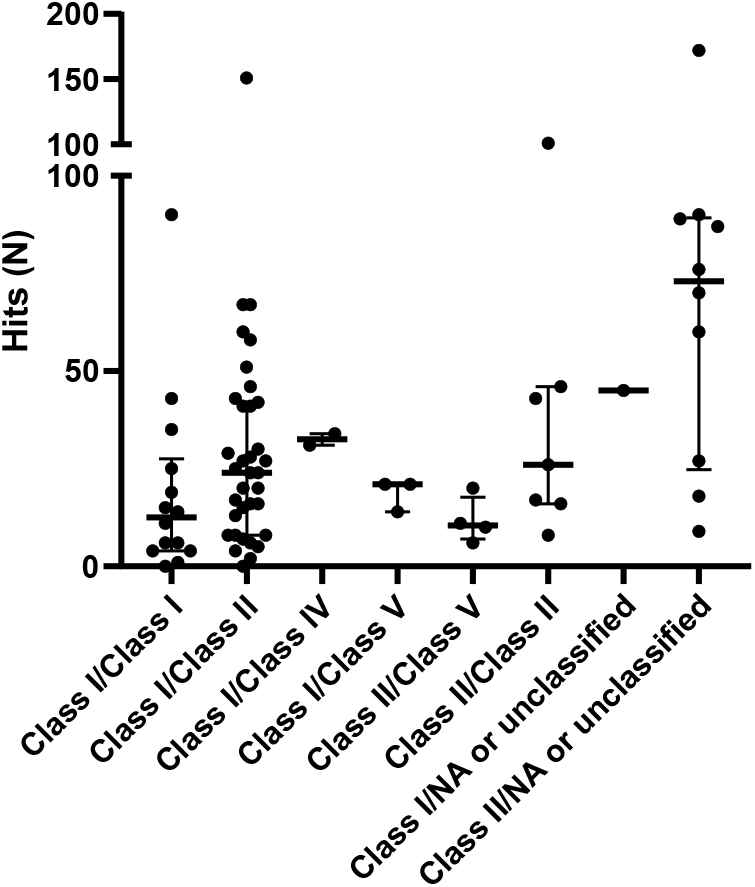
Number of hits in the primary screen for each mutation class. Medians and interquarle ranges are indicated by stripes and errorbars and are based on one technical replicate derived of one biological replicate.

**Sup. Fig. 2.**
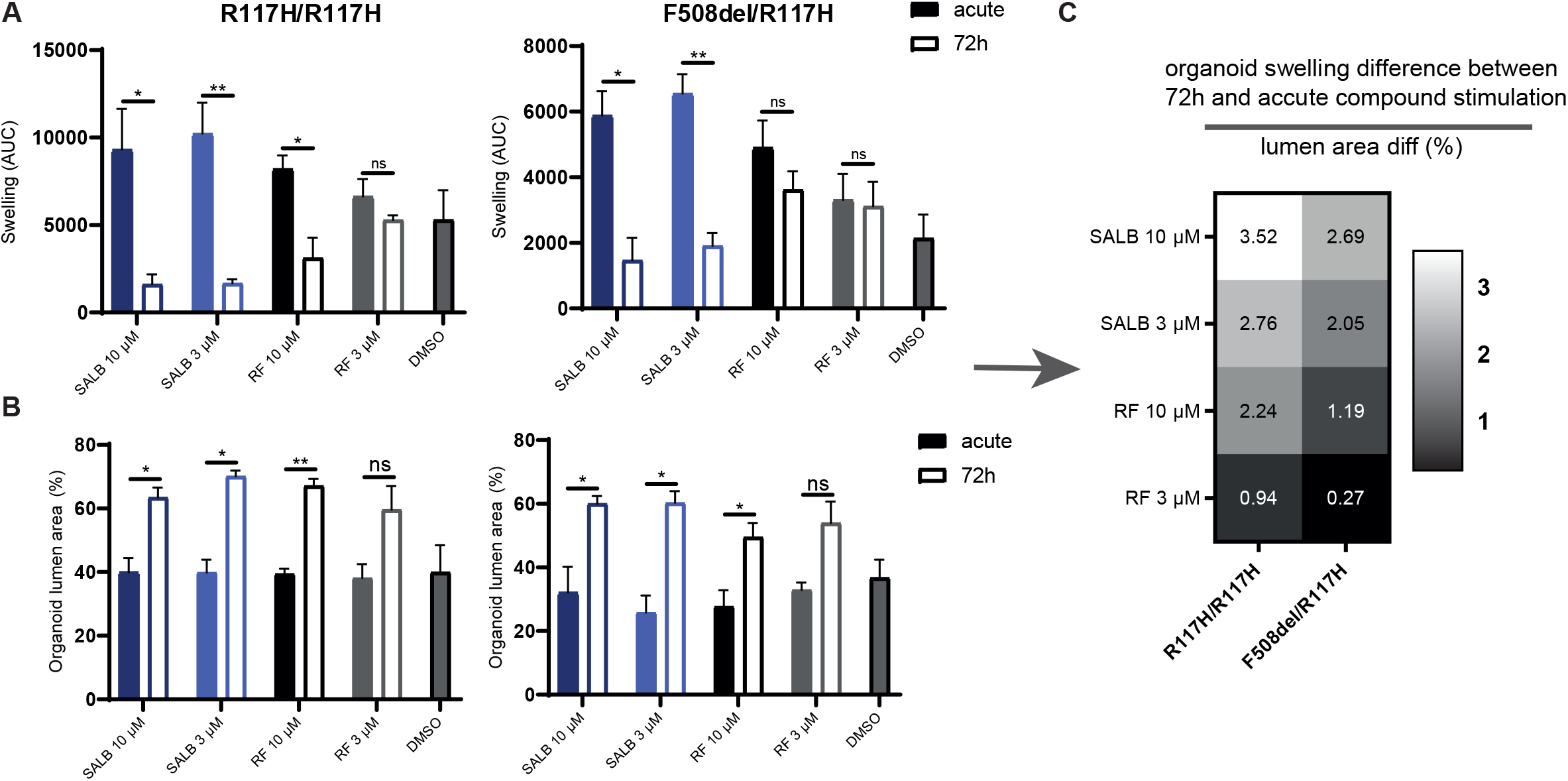
Pre-incubation with roflumilast and salbutamol results in forskolin-indepedent swelling if residual CFTR function is present. **(A)** PDIO swelling for two PDIOs upon treatment with salbutamol and roflumilast at diffierent incubations, preincubated for 72 hours or added acutely. Bars indicate the mean of three technical replicates, derived of three biological replicates. **(B)** PDIO lumen size for the PDIOs corresponding to A, upon treatment with salbutamol and roflumilast at diffierent incubations, preincubated for 72 hours or added acutely. Bars indicate the mean of three technical replicates, derived of three biological replicates. **(C)** Calculation of diffierences of organoid swelling (AUC) in response to 72hr prestimulation and acute compound treatment, divided by the lumen area (%) prior to FIS measurements. Data is shown for two PDIOs and four compounds, a value over 1 indicating that the decrease in AUC between 72hr and acute stimulation is larger than expected based on the increase of the lumen area.

**Sup. Fig. 3.**
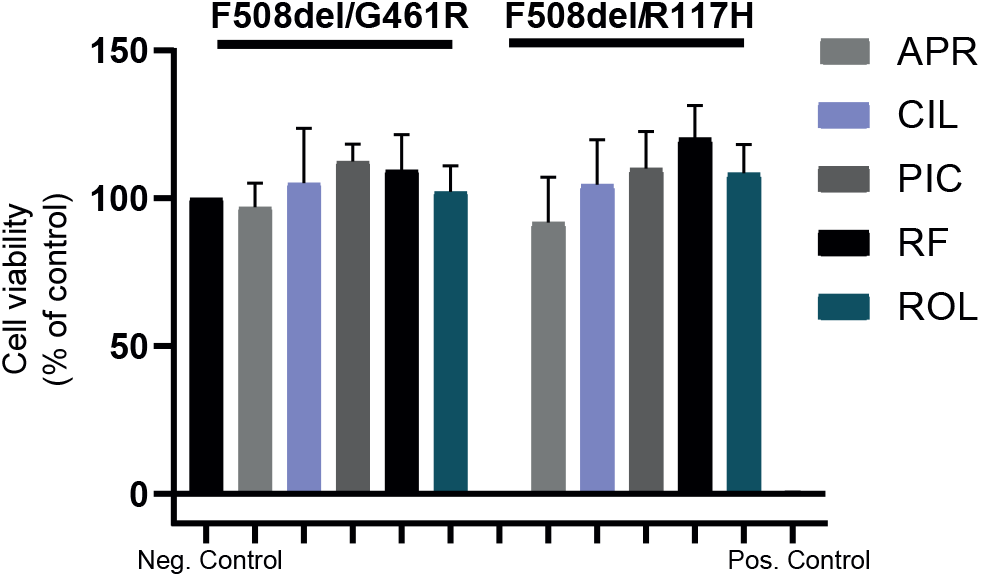
Viability of PDIOs treated with the diffierent PDE4 inhibitors. Viability was normalized to vehicle-treated negative controls and and 10% DMSO treated positive controls PDIOs. Bars indicate the mean of three technical replicates, derived of three biological replicates, with errorbars indicating the SEM.

**Sup. Fig. 4.**
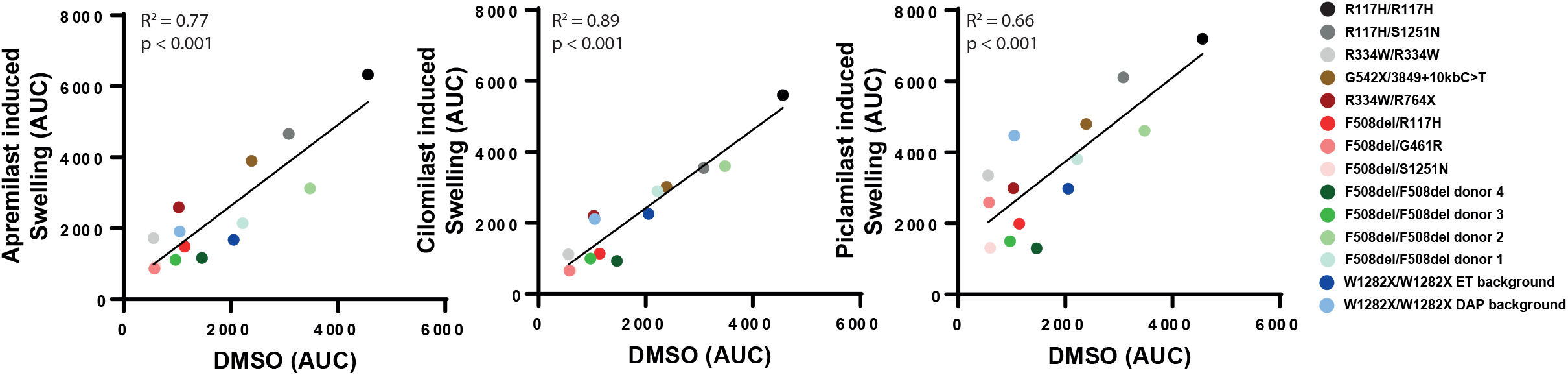
Cilomilast, Apremilast or Piclamliast induced swelling correlates with residual CFTR function. Cilomilast (left), Apremilast (middle) or Piclamliast (right) induced swelling (AUC) versus background (DMSO/other) induced swelling (AUC), for 14 PDIOs indicated by the diffierent colored dots. Dots indicate the mean of three technical replicates, derived of three biological replicates.

**Sup. Fig. 5.**
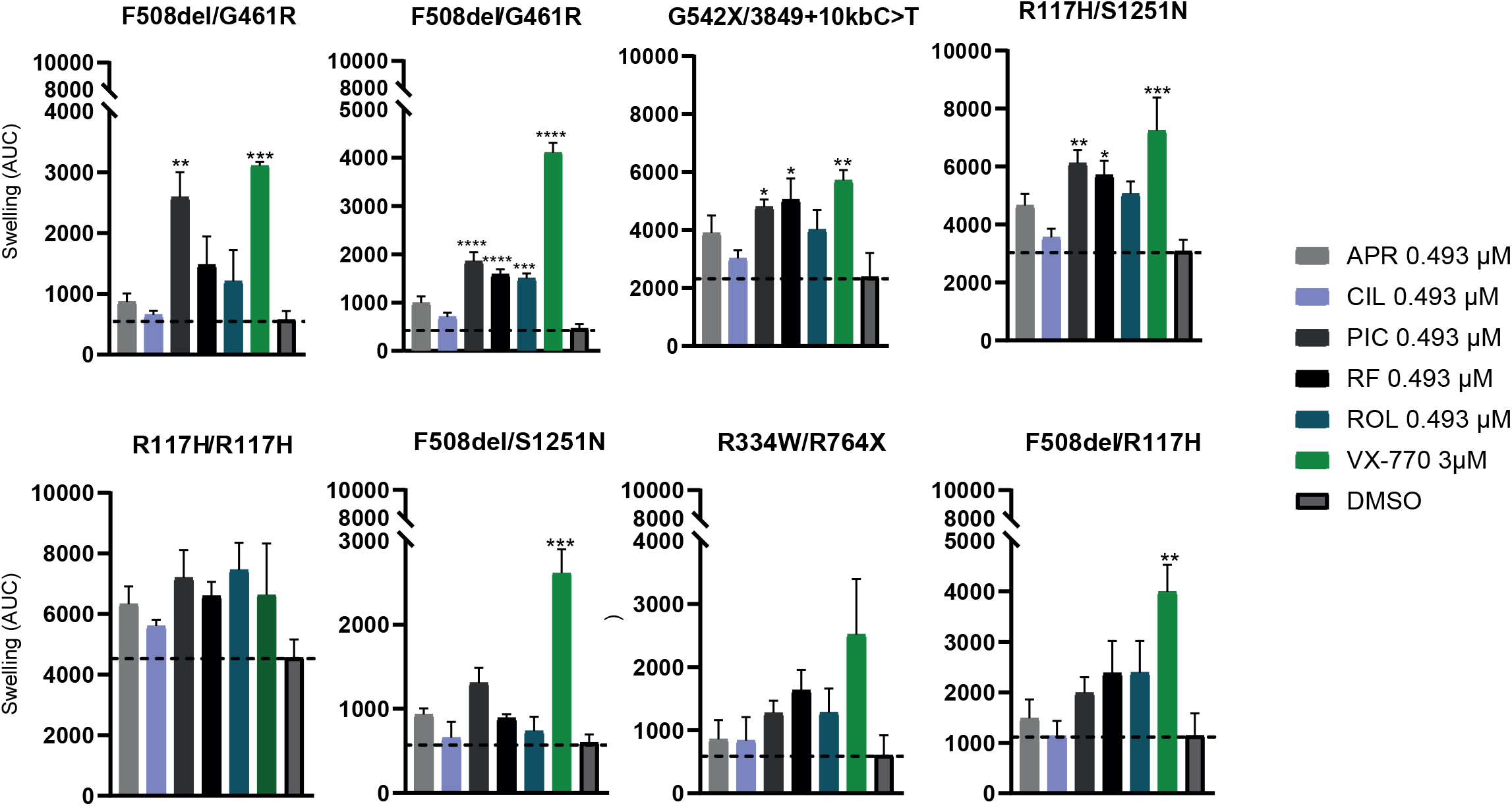
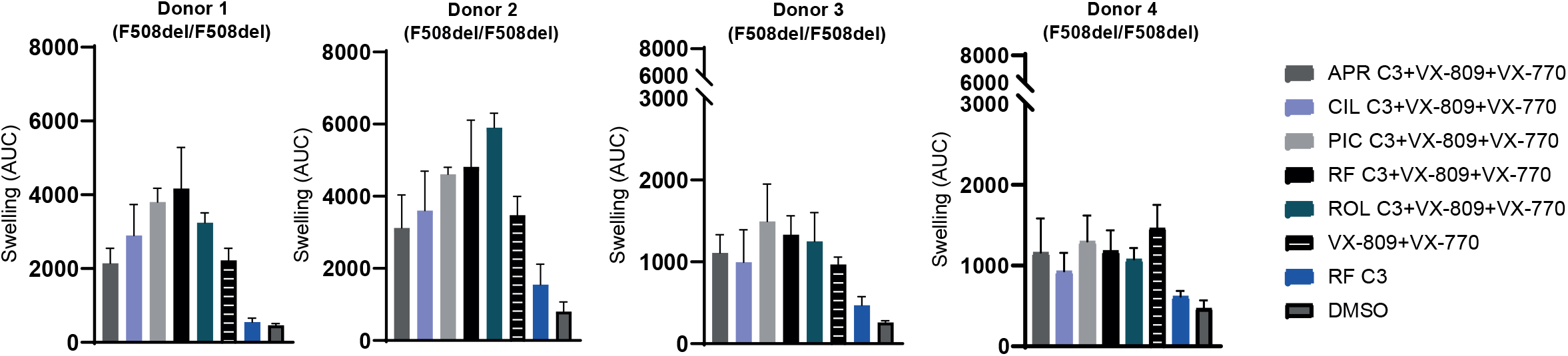
Compound-induced swelling of PDIOs with residual function and F508del/F508del-CFTR PDIOs. **(A)** PDIO swelling (AUC) of 8 PDIOs upon treatment with the various PDE4 inhibitors. Bars indicate the mean of three technical replicates, derived of three biological replicates, with errorbars indicating the SEM. **(B)** PDIO swelling (AUC) of 4 F508del/F508del PDIOs upon treatment with the various PDE4 inhibitors. Bars indicate the mean of three technical replicates, derived of three biological replicates, with errorbars indicating the SEM.

## REFERENCES

1. Ashburn, T. T. & Thor, K. B. Drug repositioning: identifying and developing new uses for existing drugs. Nat. Rev. Drug Discov. 3, 673–683 (2004).

2. Huang, F. et al. Identification of amitriptyline HCl, flavin adenine dinucleotide, azacitidine and calcitriol as repurposing drugs for influenza A H5N1 virus-induced lung injury. PLoS Pathog. 16, e1008341 (2020).

3. Despotes, K. A. & Donaldson, S. H. Current state of CFTR modulators for treatment of Cystic Fibrosis. Curr. Opin. Pharmacol. 65, 102239 (2022).

4. Vonk, A. M. et al. Protocol for Application, Standardization and Validation of the Forskolin-Induced Swelling Assay in Cystic Fibrosis Human Colon Organoids. STAR Protoc. 1, 100019 (2020).

5. Dekkers, J. F. et al. A functional CFTR assay using primary cystic fibrosis intestinal organoids. Nat. Med. 19, 939–945 (2013).

6. Dekkers, J. F. et al. Characterizing responses to CFTR-modulating drugs using rectal organoids derived from subjects with cystic fibrosis. Sci. Transl. Med. 8, 344ra84–344ra84 (2016).

7. Berkers, G. et al. Rectal organoids enable personalized treatment of cystic fibrosis. Cell Rep. 26, 1701– 1708 (2019).

8. Muilwijk, D. et al. Forskolin-induced Organoid Swelling is Associated with Long-term CF Disease Progression. Eur. Respir. J. (2022).

9. Spelier, S. et al. High-Throughput Functional Assay in Cystic Fibrosis Patient-Derived Organoids Allows Drug Repurposing. bioRxiv (2022).

10. Garnock-Jones, K. P. Roflumilast: A Review in COPD. Drugs 75, 1645–1656 (2015).

11. Shishido, H., Yoon, J. S., Yang, Z. & Skach, W. R. CFTR trafficking mutations disrupt cotranslational protein folding by targeting biosynthetic intermediates. Nat. Commun. 11, 1–11 (2020).

12. Chai, S. C., Goktug, A. N. & Chen, T. Assay validation in high throughput screening–from concept to application. Drug Discov. Dev. Mol. Med (2015).

13. Pottier, C. et al. Tyrosine kinase inhibitors in cancer: breakthrough and challenges of targeted therapy. Cancers (Basel). 12, 731 (2020).

14. Keravis, T. & Lugnier, C. Cyclic nucleotide phosphodiesterase (PDE) isozymes as targets of the intracellular signalling network: benefits of PDE inhibitors in various diseases and perspectives for future therapeutic developments. Br. J. Pharmacol. 165, 1288–1305 (2012).

15. Brewington, J. J. et al. Chronic β2AR stimulation limits CFTR activation in human airway epithelia. JCI insight 3, (2018).

16. Trzaska, C. et al. 2, 6-Diaminopurine as a highly potent corrector of UGA nonsense mutations. Nat. Commun. 11, 1–12 (2020).

17. Driehuis, E. et al. Oral mucosal organoids as a potential platform for personalized cancer therapy. Cancer Discov. (2019).

18. Folkesson, E. et al. High-throughput screening reveals higher synergistic effect of MEK inhibitor combinations in colon cancer spheroids. Sci. Rep. 10, 1–14 (2020).

19. Schütte, M. et al. Molecular dissection of colorectal cancer in pre-clinical models identifies biomarkers predicting sensitivity to EGFR inhibitors. Nat. Commun. 8, 1–19 (2017).

20. Jiang, J. X. et al. A new platform for high-throughput therapy testing on iPSC-derived lung progenitor cells from cystic fibrosis patients. Stem cell reports 16, 2825–2837 (2021).

21. Berg, A. et al. High-throughput surface liquid absorption and secretion assays to identify F508del CFTR correctors using patient primary airway epithelial cultures. SLAS Discov. Adv. Life Sci. R&D 24, 724–737 (2019).

22. Vijftigschild, L. A. W. et al. 2-Adrenergic receptor agonists activate CFTR in intestinal organoids and subjects with cystic fibrosis. European Respiratory Journal vol. 48 768–779 at https://doi.org/10.1183/13993003.01661-2015 (2016).

23. Medeiros, B. C., Possick, J. & Fradley, M. Cardiovascular, pulmonary, and metabolic toxicities complicating tyrosine kinase inhibitor therapy in chronic myeloid leukemia: Strategies for monitoring, detecting, and managing. Blood Rev. 32, 289–299 (2018).

24. Mingard, C., Paech, F., Bouitbir, J. & Krähenbühl, S. Mechanisms of toxicity associated with six tyrosine kinase inhibitors in human hepatocyte cell lines. J. Appl. Toxicol. 38, 418–431 (2018).

25. Cazzola, M., Calzetta, L., Rogliani, P. & Matera, M. G. The discovery of roflumilast for the treatment of chronic obstructive pulmonary disease. Expert Opin. Drug Discov. 11, 733–744 (2016).

26. Zhao, Y. U., Zhang, H.-T. & O’Donnell, J. M. Inhibitor binding to type 4 phosphodiesterase (PDE4) assessed using [3H] piclamilast and [3H] rolipram. J. Pharmacol. Exp. Ther. 305, 565–572 (2003).

27. Huai, Q. et al. Enantiomer discrimination illustrated by the high resolution crystal structures of type 4 phosphodiesterase. J. Med. Chem. 49, 1867–1873 (2006).

28. Schafer, P. H. et al. Apremilast, a cAMP phosphodiesterase-4 inhibitor, demonstrates anti-inflammatory activity in vitro and in a model of psoriasis. Br. J. Pharmacol. 159, 842–855 (2010).

29. Yokota, H. et al. Relationship between Plasma Concentrations of Afatinib and the Onset of Diarrhea in Patients with Non-Small Cell Lung Cancer. Biology (Basel). 10, 1054 (2021).

30. Duan, T., Cil, O., Thiagarajah, J. R. & Verkman, A. S. Intestinal epithelial potassium channels and CFTR chloride channels activated in ErbB tyrosine kinase inhibitor diarrhea. JCI insight 4, (2019).

31. Reilly, R. et al. Targeting the PI3K/Akt/mTOR signalling pathway in Cystic Fibrosis. Sci. Rep. 7, 1–13 (2017).

32. Verkman, A. S. & Galietta, L. J. Chloride channels as drug targets. Nat Rev Drug Discov. 8, 153–171 (2009).

33. Ma, X. et al. Activation of GABAA receptors in colon epithelium exacerbates acute colitis. Front. Immunol. 9, 1–18 (2018).

34. Li, Y., Xiang, Y. Y., Lu, W. Y., Liu, C. & Li, J. A novel role of intestine epithelial GABAergic signaling in regulating intestinal fluid secretion. Am. J. Physiol. - Gastrointest. Liver Physiol. 303, 453–460 (2012).

35. Hyland, N. P. & Cryan, J. F. A gut feeling about GABA: Focus on GABAB receptors. Front. Pharmacol. OCT, 1–9 (2010).

36. Amatngalim, G. D. et al. Measuring cystic fibrosis drug responses in organoids derived from 2D differentiated nasal epithelia. Life Sci. Alliance 5, (2022).

37. Flume, P. A. et al. Ivacaftor in subjects with cystic fibrosis who are homozygous for the F508del-CFTR mutation. Chest 142, 718–724 (2012).

38. Rowe, S. M. et al. Lumacaftor/ivacaftor treatment of patients with cystic fibrosis heterozygous for F508del-CFTR. Ann. Am. Thorac. Soc. 14, 213–219 (2017).

39. Rowe, S. M. et al. Tezacaftor–Ivacaftor in Residual-Function Heterozygotes with Cystic Fibrosis. N. Engl. J. Med. 377, 2024–2035 (2017).

40. Wainwright, C. E. et al. Lumacaftor–Ivacaftor in Patients with Cystic Fibrosis Homozygous for Phe508del CFTR. N. Engl. J. Med. 373, 220–231 (2015).

41. Moss, R. B. et al. Efficacy and safety of ivacaftor in patients with cystic fibrosis who have an Arg117His-CFTR mutation: A double-blind, randomised controlled trial. Lancet Respir. Med. 3, 524–533 (2015).

42. K., D. B. et al. Efficacy and safety of ivacaftor in patients with cystic fibrosis and a non-G551D gating mutation. J. Cyst. Fibros. 13, 674–680 (2014).

