## Supplemental Data for "FDA-Approved Drug Screening in Patient-Derived Organoids Demonstrates Potential of Drug Repurposing for Rare Cystic Fibrosis Genotypes"

Sup. Fig. 1

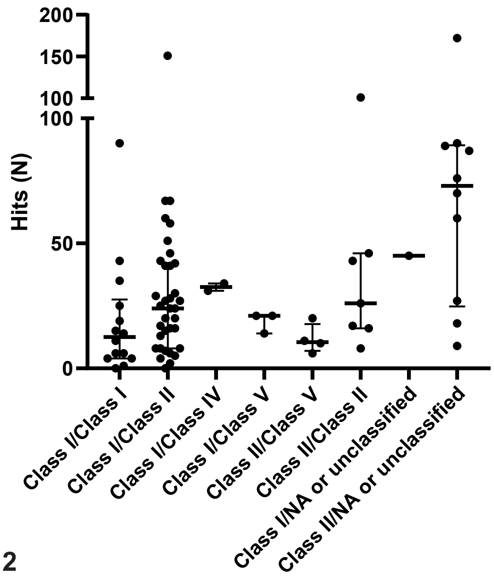


Sup. Fig. 2

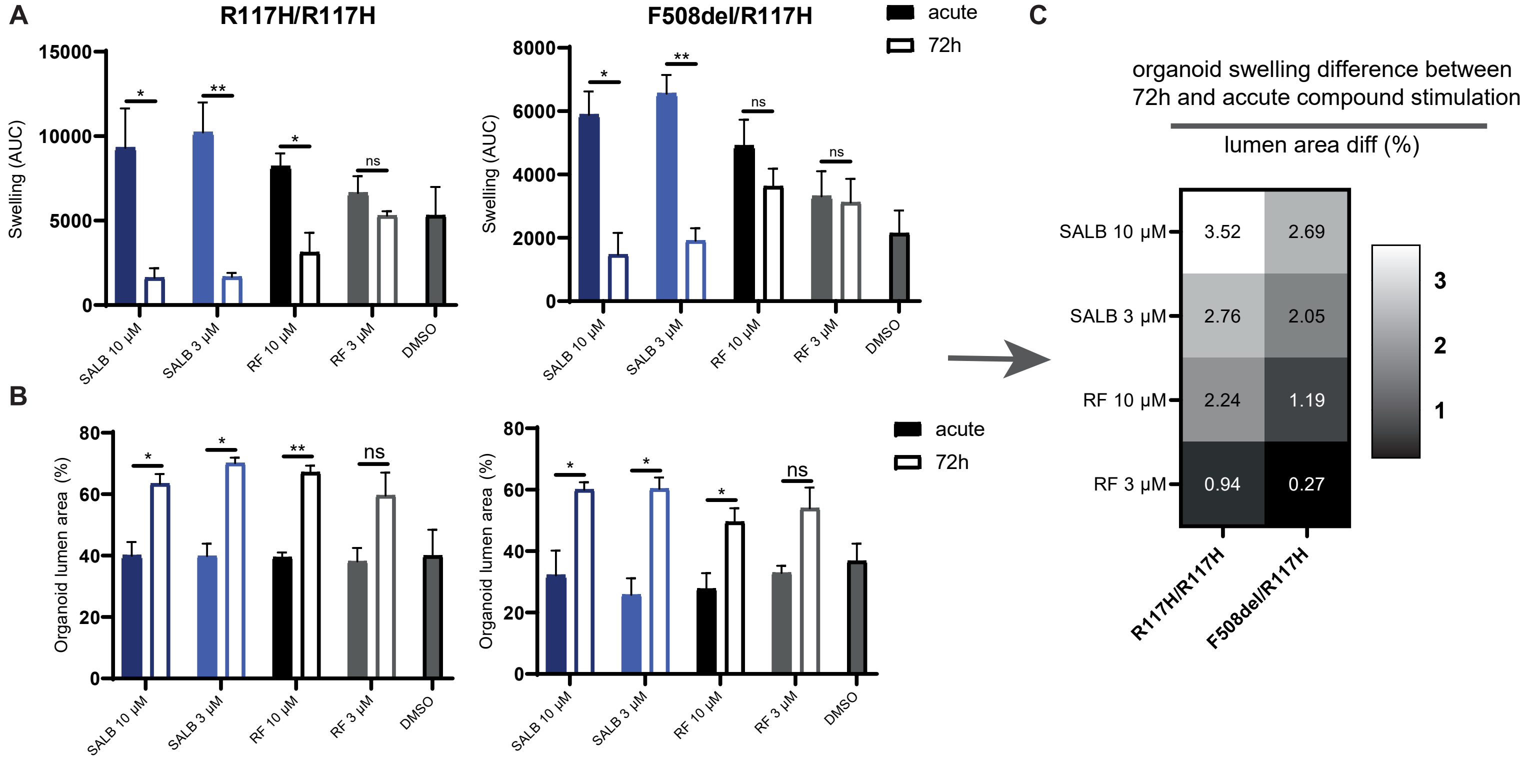


Sup. Fig. 3

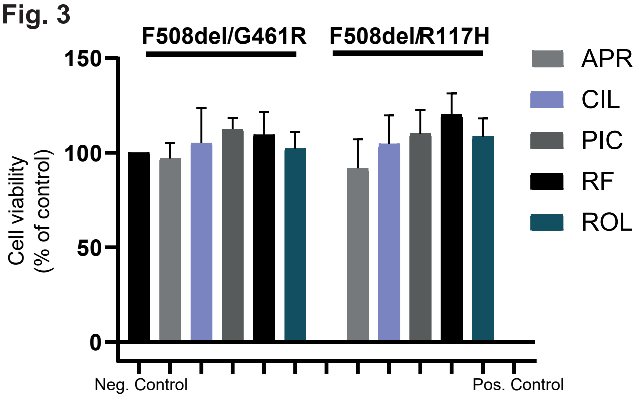


Sup. Fig. 4

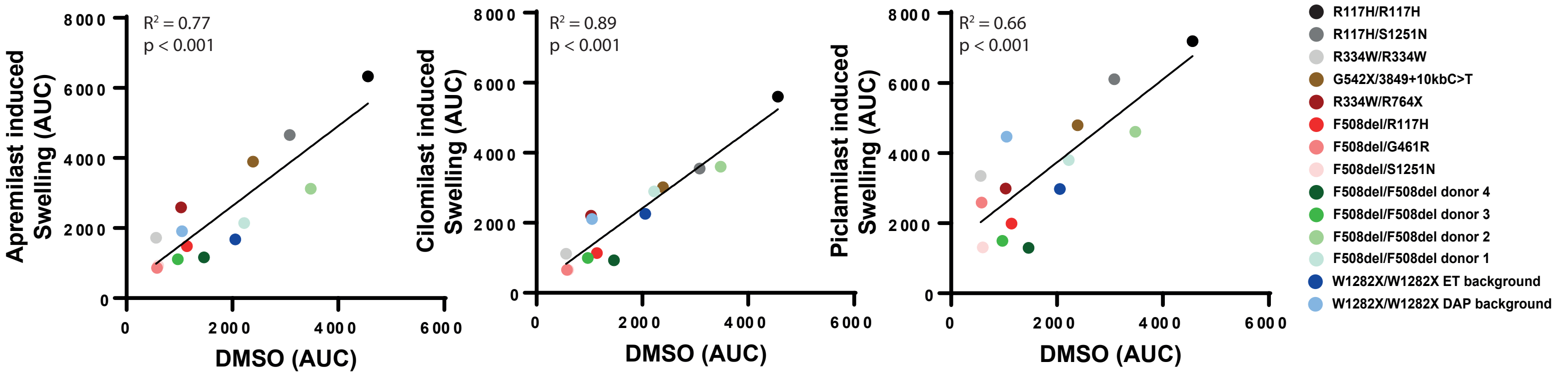

Sup. Fig. 5A

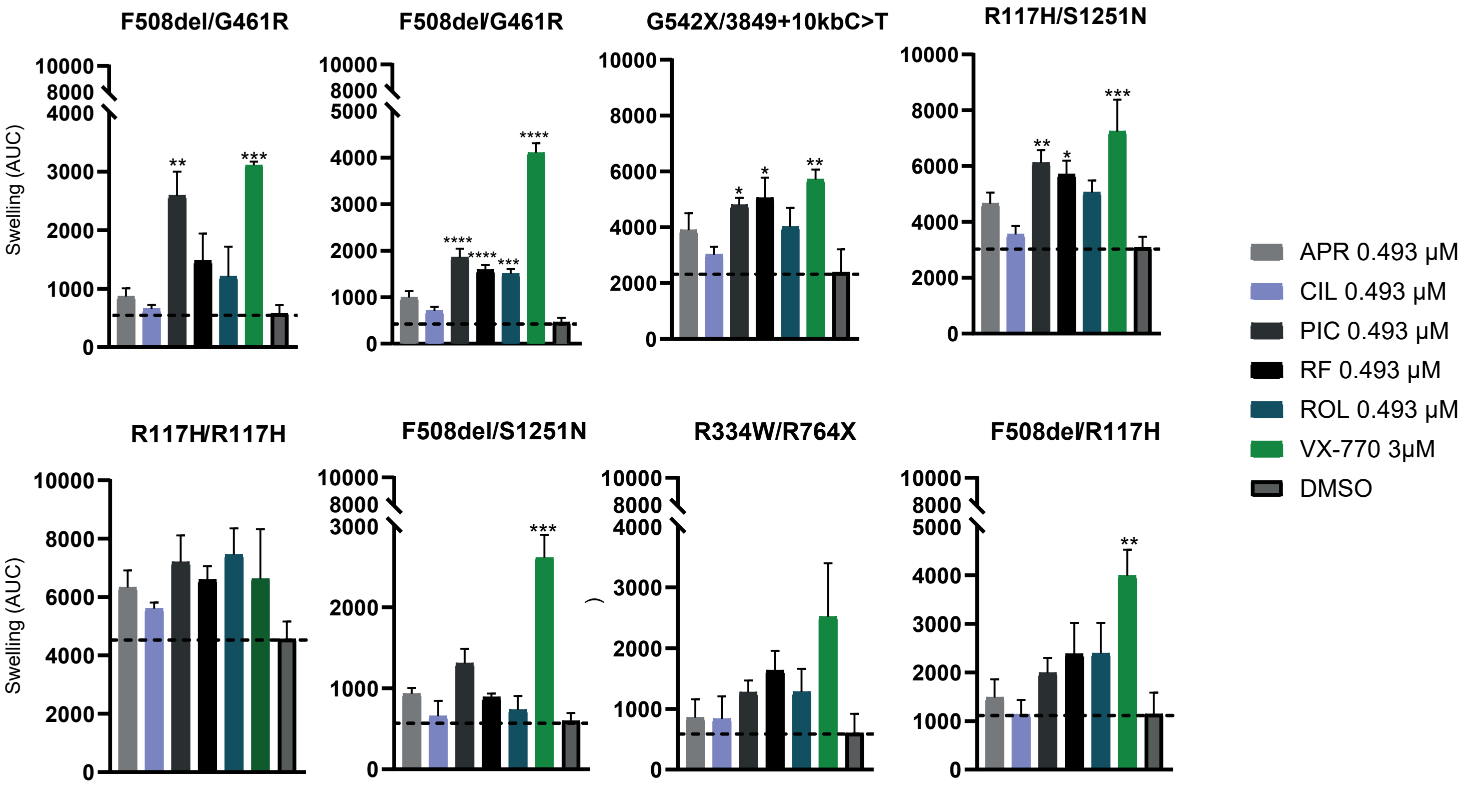


Sup. Fig. 5B

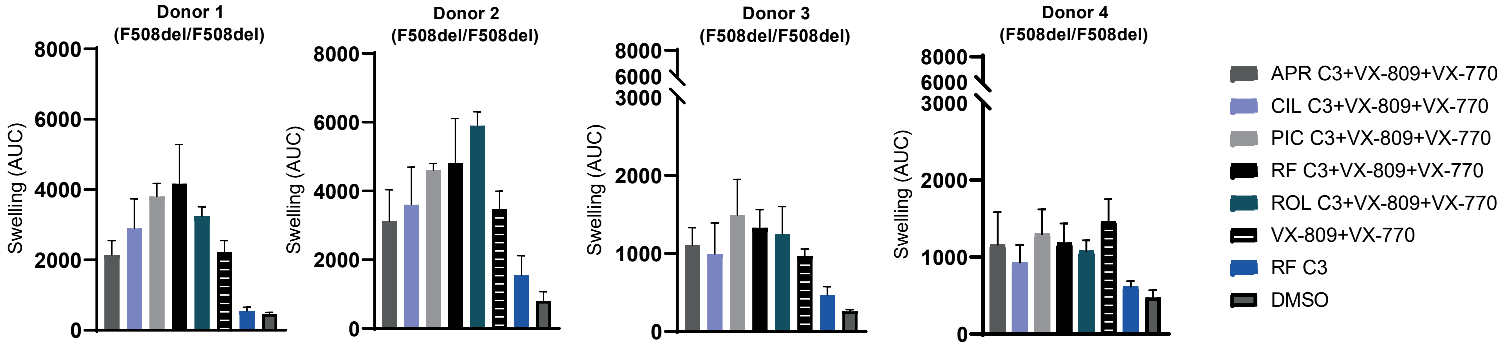


**Sup. Fig. 1 Number of hits in the primary screen for each mutation class.**
Medians and interquartile ranges are indicated by stripes and errorbars and are based on one technical replicate derived of one biological replicate.

**Sup. Fig. 2 Pre-incubation with roflumilast and salbutamol results in forskolin-indepedent swelling if residual CFTR function is present**
**(A)** PDIO swelling for two PDIOs upon treatment with salbutamol and roflumilast at different incubations, preincubated for 72 hours or added acutely. Bars indicate the mean of three technical replicates, derived of three biological replicates. **(B)** PDIO lumen size for the PDIOs corresponding to A, upon treatment with salbutamol and roflumilast at different incubations, preincubated for 72 hours or added acutely. Bars indicate the mean of three technical replicates, derived of three biological replicates. **(C)** Calculation of differences of organoid swelling (AUC) in response to 72hr prestimulation and acute compound treatment, divided by the lumen area (%) prior to FIS measurements. Data is shown for two PDIOs and four compounds, a value over 1 indicating that the decrease in AUC between 72hr and acute stimulation is larger than expected based on the increase of the lumen area.

**Sup. Fig. 3 Viability of PDIOs treated with the different PDE4 inhibitors**
Viability was normalized to vehicle-treated negative controls and and 10% DMSO treated positive controls PDIOs. Bars indicate the mean of three technical replicates, derived of three biological replicates, with errorbars indicating the SEM.

**Sup. Fig. 4 Cilomilast, Apremilast or Piclamliast induced swelling correlates with residual CFTR function**
Cilomilast (left), Apremilast (middle) or Piclamliast (right) induced swelling (AUC) versus background (DMSO/other) induced swelling (AUC), for 14 PDIOs indicated by the different colored dots. Dots indicate the mean of three technical replicates, derived of three biological replicates.

**Sup. Fig. 5 Compound-induced swelling of PDIOs with residual function and F508del/F508del-CFTR PDIOs**
**(A)** PDIO swelling (AUC) of 8 PDIOs upon treatment with the various PDE4 inhibitors. Bars indicate the mean of three technical replicates, derived of three biological replicates, with errorbars indicating the SEM. **(B)** PDIO swelling (AUC) of 4 F508del/F508del PDIOs upon treatment with the various PDE4 inhibitors. Bars indicate the mean of three technical replicates, derived of three biological replicates, with errorbars indicating the SEM.

**Supplemental Table 1**

Overview genotypes and corresponding CFTR classes of PDIOs included in primary screen, according to reference in table. Mutation categories were summarized by A: Class I/Class I, B: Class I/Class II, C: Class I/Class IV, D: Class I/Class V, E: Class II/Class V, F: Class II/Class II, G: Class I/NA or Unclassified, H: Class II/NA or Unclassified

| Donor number | Genotype | Forskolin (µM) | Allele 1 Class | Allele 2 Class | Reference Allele Class | Mutation category |
|---|---|---|---|---|---|---|
| donor 01 | 1677delTA/3120+1G>A | 5 | I | I | Muijlwijk (2022) | A |
| donor 02 | 1717-1G>A/2183AA>G | 5 | I | I | Muijlwijk ((2022)) | A |
| donor 03 | 1717-1G>T/3905insT | 5 | I | I | Muijlwijk (2022) | A |
| donor 04 | 1811+1G>C/1811+1G>C | 5 | I | I | Muijlwijk (2022) | A |
| donor 05 | 1811+1G>C/1811+1G>C | 5 | I | I | Muijlwijk (2022) | A |
| donor 06 | 1811+1G>C/1811+1G>C | 5 | I | I | Muijlwijk (2022) | A |
| donor 07 | 1811+1G>C/1811+1G>C | 5 | I | I | Muijlwijk (2022) | A |
| donor 08 | 2789+5G>T/711+1G>T | 0.128 | V | I | Muijlwijk (2022) | D |
| donor 09 | 3272-26A->G/G970R | 0.128 | V | I | Muijlwijk (2022) | D |
| donor 10 | 579+1G>T/CFTRdele11 | 5 | I | I | Muijlwijk (2022) | A |
| donor 11 | 711+1G>T/711+1G>T | 5 | I | I | Muijlwijk (2022) | A |
| donor 12 | A455E/1343delG | 0.8 | II | I | - | B |
| donor 13 | A455E/E60X | 0.128 | II | I | Muijlwijk (2022) | B |
| donor 14 | A455E/NA | 0.128 | II | NA | Muijlwijk (2022) | H |
| donor 15 | A46D/A46D | 0.8 | II | II | Muijlwijk (2022) | F |
| donor 16 | E60X/4015delATTT | 5 | I | I | - | A |
| donor 17 | E92K/E92K | 0.128 | II | II | Muijlwijk (2022) | F |
| donor 18 | F508del/(TG)13(T)5 | 0.128 | II | V | Muijlwijk (2022) | E |
| donor 19 | F508del/(TG)13(T)5 | 5 | II | V | Muijlwijk (2022) | E |
| donor 20 | F508del/1078delT | 5 | II | I | Muijlwijk (2022) | B |
| donor 21 | F508del/1294del7 | 5 | II | I | Niet in CFTR2 | B |
| donor 22 | F508del/1813insC | 5 | II | I | Muijlwijk (2022) | B |
| donor 23 | F508del/2143delT | 5 | II | I | - | B |
| donor 24 | F508del/2143delT | 5 | II | I | - | B |
| donor 25 | F508del/2183AA>G | 0.8 | II | I | Muijlwijk (2022) | B |
| donor 26 | F508del/2184delA | 5 | II | I | Muijlwijk (2022) | B |
| donor 27 | F508del/3272-26A>G | 0.128 | II | V | Muijlwijk (2022) | E |
| donor 28 | F508del/3500-2A->G | 5 | II | I | - | B |
| donor 29 | F508del/365-366insT(W79fs) | 0.8 | II | I | Muijlwijk (2022) | B |
| donor 30 | F508del/3659delC | 5 | II | I | Muijlwijk (2022) | B |
| donor 31 | F508del/394delTT | 0.8 | II | I | Muijlwijk (2022) | B |
| donor 32 | F508del/4374+1G>T | 5 | II | I | Muijlwijk (2022) | B |
| donor 33 | F508del/4382delA | 5 | II | V | Muijlwijk (2022) | E |

| | | | | | | |
|---|---|---|---|---|---|---|
| donor 34 | F508del/711+1G>T | 5 | II | I | Muijlwijk (2022) | B |
| donor 35 | F508del/711+1G>T | 5 | II | I | Muijlwijk (2022) | B |
| donor 36 | F508del/711+1G>T | 5 | II | I | Muijlwijk (2022) | B |
| donor 37 | F508del/CFTRdele17a.17b | 5 | II | I | Muijlwijk (2022) | B |
| donor 38 | F508del/CFTRdele17a.17b | 5 | II | I | Muijlwijk (2022) | B |
| donor 39 | F508del/CFTRdele19.20 | 5 | II | I | Muijlwijk (2022) | B |
| donor 40 | F508del/CFTRdele2.3 | 5 | II | I | - | B |
| donor 41 | F508del/CFTRdele2.3 | 5 | II | I | - | B |
| donor 42 | F508del/E60X | 5 | II | I | Muijlwijk (2022) | B |
| donor 43 | F508del/E730X | 0.8 | II | I | Muijlwijk (2022) | B |
| donor 44 | F508del/G1249R | 5 | II | Unclassified | Muijlwijk (2022) | H |
| donor 45 | F508del/G461R | 0.128 | II | Unclassified | Muijlwijk (2022) | H |
| donor 46 | F508del/G461R | 0.128 | II | Unclassified | Muijlwijk (2022) | H |
| donor 47 | F508del/G461R | 0.128 | II | Unclassified | Muijlwijk (2022) | H |
| donor 48 | F508del/G550X | 5 | II | I | Muijlwijk (2022) | B |
| donor 49 | F508del/G550X | 5 | II | I | Muijlwijk (2022) | B |
| donor 50 | F508del/4046delG | 0.128 | II | Unclassified | Muijlwijk (2022) | H |
| donor 51 | F508del/L927P | 5 | II | Unclassified | Muijlwijk (2022) | H |
| donor 52 | F508del/L927P | 5 | II | Unclassified | Muijlwijk (2022) | H |
| donor 53 | F508del/R1066C | 0.8 | II | II | Muijlwijk (2022) | F |
| donor 54 | F508del/R347P | 0.128 | II | II | Muijlwijk (2022) | F |
| donor 55 | F508del/R347P | 0.128 | II | II | Muijlwijk (2022) | F |
| donor 56 | F508del/R347P | 0.128 | II | II | Muijlwijk (2022) | F |
| donor 57 | F508del/R74P | 0.8 | II | Unclassified | Muijlwijk (2022) | H |
| donor 58 | F508del/S489X | 5 | II | I | Muijlwijk (2022) | B |
| donor 59 | F508del/W1282X | 5 | II | I | Muijlwijk (2022) | B |
| donor 60 | F508del/W1282X | 5 | II | I | Muijlwijk (2022) | B |
| donor 61 | F508del/Y1092X | 5 | II | I | Muijlwijk (2022) | B |
| donor 62 | F508del/Y1092X | 0.8 | II | I | Muijlwijk (2022) | B |
| donor 63 | F508del/Y1092X | 5 | II | I | Muijlwijk (2022) | B |
| donor 64 | F508del/Y109D | 0.8 | II | Unclassified | Muijlwijk (2022) | H |
| donor 65 | G542X/CFTRdele2.3 | 5 | I | I | - | A |
| donor 66 | G542X/G542X | 5 | I | I | Muijlwijk (2022) | A |
| donor 67 | G542X/R1066C | 5 | I | II | - | B |
| donor 68 | N1303K/G550X | 5 | II | I | Muijlwijk (2022) | B |
| donor 69 | R1162X/3539del16 | 5 | I | I | - | A |

| | | | | | | |
|---|---|---|---|---|---|---|
| donor 70 | R1162X/3659delC | 5 | I | I | Muijlwijk (2022) | A |
| donor 71 | R117H-7T/T1857delT | 0.128 | IV | I | - | C |
| donor 72 | R334W/R334W | 0.128 | II | II | Muijlwijk (2022) | F |
| donor 73 | R334W/R764X | 0.128 | IV | I | Muijlwijk (2022) | C |
| donor 74 | R553X/NA | 0.128 | I | NA | Muijlwijk (2022) | G |
| donor 75 | Y275X/A559T | 0.8 | I | II | Shishido, 2020 | B |
| donor 76 | Y849X/2789+5G>A | 0.128 | I | V | Muijlwijk (2022) | D |

**Supplemental Table 2**
Overview of hits in primary FDA screen per mutation category, including mean number of hits, the top-responders and the lowest responders.

| Category | Median number of hits | Hits of Top Responders (N) | Hits of Top Non-Responders (N) |
|---|---|---|---|
| Class I/Class I | 12.5 | 90 R1162X/3659delC donor 70<br>43 579+1G>T/CFTRdele11 donor 10<br>35 711+1G>T/711+1G>T donor 11 | 4 R1162X/3539del16 donor 69<br>1 1811+1G>C/1811+1G>C donor 04<br>0 1811+1G>C/1811+1G>C donor 06 |
| Class I/Class II | 24 | 151 F508del/394delTT donor 31<br>67 F508del/Y1092X donor 63<br>67 F508del/CFTRdele19.20 donor 39 | 4 F508del/3500-2A->G donor 28<br>2 F508del/711+1G>T donor 35<br>0 F508del/711+1G>T donor 34 |
| Class I/Class IV | 32.5 | 34 R117H-7T/T1857delT donor 71 | 31 R334W/R764X donor 73 |
| Class I/Class V | 21 | 21 3272-26A->G/G970R donor 09<br>21 Y849X/2789+5G>A donor 76 | 14 2789+5G>T/711+1G>T donor 08 |
| Class II/Class V | 10.5 | 11 F508del/(TG)13(T)5 donor 18<br>20 F508del/(TG)13(T)5 donor 19 | 10 F508del/3272-26A>G donor 27<br>6 F508del/4382delA donor 33 |
| Class II/Class II | 26 | 101 F508del/R347P donor 54<br>46 F508del/R347P donor 56<br>43 F508del/R1066C donor 53 | 17 E92K/E92K donor 17<br>16 A46D/A46D donor 15<br>8 R334W/R334W donor 72 |
| Class I/NA or unclassified | 45 | 45 R553X/NA donor 74 | |
| Class II/NA or unclassified | 73 | 172 F508del/G461R donor 47<br>90 F508del/L927P donor 52<br>89 F508del/L927P donor 51 | 27 F508del/G1249R donor 44<br>18 F508del/R74P donor 57<br>9 F508del/Y109D donor 64 |

**Supplemental Table 3**
Overview of genotypes PDIOs included in the Roflumilast screen, corresponding to **Figure 4E and 4F**.

| CF-number | Genotype | fsk concentration (µM) |
|---|---|---|
| CF0021 | F508del/(TG)13(T)5 | 0.128 |
| CF0024 | F508del/(TG)13(T)5 | 0.128 |
| CF0025 | F508del/(TG)13(T)5 | 0.128 |
| CF0055 | F508del/G461R | 0.128 |
| CF0063 | R117H-7T/T1857delT | 0.128 |
| CF00160 | A455E/E60X | 0.128 |
| CF00175 | F508del/365-366insT(W79fs) | 0.128 |
| CF0176 | F508del/R347P | 0.128 |
| CF0198 | R553X/NA | 0.128 |
| CF0215 | F508del/G628R | 0.128 |
| CF0227 | 3272-26A->G/G970R | 0.128 |
| CF0228 | F508del/3272-26A>G | 0.128 |
| CF0238 | F508del/R347P | 0.128 |
| CF0284 | R117H-7T/UNK | 0.128 |
| CF0333 | UNK/UNK | 0.128 |
| CF0349 | F508del/4382delA | 0.128 |
| CF0377 | F508del/UNK | 0.128 |
| CF0401 | 3849+10kbC>T/R347H | 0.128 |
| CF0406 | F508del/4382delA | 0.128 |
| CF0419 | A455E/1343delG | 0.128 |

| | | |
|---|---|---|
| CF0470 | A455E/UNK | 0.128 |
| CF0478 | 3905insT/D1152H | 0.128 |
| CF0480 | F508del/R75Q | 0.128 |
| CF0497 | F508del/3849+10kbC>T | 0.128 |
| CF0534 | F508del/I1027T | 0.128 |
| CF0171 | F508del/Gly1349fs | 0.8 |
| CF0219 | F508del/R347P | 0.8 |
| CF0225 | F508del/L1034P | 0.8 |
| CF0303 | A455E/NA | 0.8 |
| CF0305 | 2789+5G>A/UNK | 0.8 |
| CF0308 | F508del/621+1G>T | 0.8 |
| CF0348 | F508del/R74P | 0.8 |
| CF0360 | A46D/A46D | 0.8 |
| CF0403 | F508del/R1066C | 0.8 |
| CF0404 | F508del/3849+10kbC>T | 0.8 |
| CF0428 | W1282X/W1282X | 0.8 |
| CF0523 | F508del/S18I | 0.8 |
| CF0008 | F508del/L927P | 5 |
| CF0012 | E60X/4015delATTT | 5 |
| CF0033 | F508del/Y1092X | 5 |
| CF0049 | F508del/2183AA>G | 5 |
| CF0050 | F508del/E60X | 5 |
| CF0068 | F508del/3500-2A->G | 5 |
| CF0075 | 1811+1G>C/1811+1G>C | 5 |
| CF0134 | F508del/711+1G>T | 5 |
| CF0135 | F508del/711+1G>T | 5 |
| CF0139 | 1677delTA/IVS16+1G>A(3120+1G>A) | 5 |
| CF0168 | 1811+1G>C/1811+1G>C | 5 |
| CF0181 | 1717-1G>A/2183AA>G | 5 |
| CF0188 | 1811+1G>C/1811+1G>C | 5 |
| CF0191 | F508del/4374+1G>T | 5 |
| CF0206 | F508del/W1282X | 5 |
| CF0217 | F508del/E730X | 5 |
| CF0224 | 711+1G>T/711+1G>T | 5 |
| CF0236 | 579+1G>T/CFTRdele11 | 5 |
| CF0239 | F508del/L927P | 5 |
| CF0248 | F508del/CFTRdele17a.17b | 5 |
| CF0250 | F508del/Y1092X | 5 |
| CF0269 | F508del/G85E | 5 |
| CF0270 | F508del/2184delA | 5 |
| CF0271 | 1811+1G>C/1811+1G>C | 5 |
| CF0276 | F508del/G550X | 5 |
| CF0282 | G542X/G542X | 5 |
| CF0289 | R1162X/3659delC | 5 |
| CF0291 | F508del/711+1G>T | 5 |
| CF0296 | N1303K/G550X | 5 |
| CF0297 | F508del/1294del7 | 5 |
| CF0304 | F508del/711+1G>T | 5 |
| CF0314 | F508del/Y1092X | 5 |
| CF0315 | F508del/Y1092X | 5 |
| CF0317 | F508del/CFTRdele19.20 | 5 |
| CF0335 | F508del/1078delT | 5 |
| CF0338 | F508del/CFTRdele2.3 | 5 |
| CF0340 | F508del/Y109D | 5 |
| CF0341 | F508del/S489X | 5 |
| CF0342 | F508del/2143delT | 5 |
| CF0346 | F508del/1813insC | 5 |
| CF0351 | F508del/E60X | 5 |
| CF0354 | F508del/L927P | 5 |

| | | |
|---|---|---|
| CF0355 | F508del/L927P | 5 |
| CF0357 | R1162X/3539del16 | 5 |
| CF0359 | F508del/394delTT | 5 |
| CF0361 | A46D/A46D | 5 |
| CF0366 | F508del/UNK | 5 |
| CF0373 | F508del/Y849X | 5 |
| CF0375 | F508del/2143delT | 5 |
| CF0378 | F508del/3905insT | 5 |
| CF0384 | F508del/CFTRdele17a.17b | 5 |
| CF0386 | G542X/CFTRdele2.3 | 5 |
| CF0388 | F508del/G550X | 5 |
| CF0391 | G542X/R1066C | 5 |
| CF0392 | W1282X/L927P | 5 |
| CF0396 | Y849X/2789+5G>A | 5 |
| CF0399 | F508del/CFTRdele2.3 | 5 |
| CF0400 | R1066C/R1066H | 5 |
| CF0412 | Y275X/A559T | 5 |
| CF0444 | F508del/G550X | 5 |
| CF0457 | F508del/3659delC | 5 |
| CF0460 | F508del/L732X | 5 |
| CF0468 | F508del/UNK | 5 |
| CF0477 | F508del/3659delC | 5 |
| CF0488 | F508del/Q493X | 5 |
| CF0489 | G542X/W679X | 5 |
| CF0496 | F508del/W1282X | 5 |
| CF0515 | F508del/W1282X | 5 |
| CF0520 | F508del/Y849X | 5 |
| CF0582 | F508del/G551D | 5 |

**Supplemental Table 4**
Overview of genotypes of PDIOs included in the CFTR modulator screen, corresponding to **Figure 5A & 5B**

| CF-number | Genotype |
|---|---|
| CF0008, CF0354, CF0239, CF0355 | F508del/L927P |
| CF0012 | E60X/4015delATTT |
| CF0031 | R117H-7T-9T/A455E |
| CF0033, CF0315, CF0314, CF0394, CF0250 | F508del/Y1092X |
| CF0035, CF0609, CF0025, CF0024, CF0021 | F508del/(TG)13(T)5 |
| CF0044 | R117H-7T/R1162X |
| CF0049 | F508del/2183AA>G |
| CF0050, CF0543, CF0351 | F508del/E60X |
| CF0055, CF0170, CF0169 | F508del/G461R |
| CF0060, CF0007, CF0011, CF0046 | F508del/A455E |
| CF0063 | R117H-7T/T1857delT |
| CF0068 | F508del/3500-2A->G |
| CF0073, CF0161 | F508del/G1249R |
| CF0077, CF0076, CF0211, CF0040, CF0212 | F508del/R1162X |
| CF0088 | A455E/S1251N |
| CF0092 | R334W/R764X |
| CF0122 | R117H/I1139V |

| | |
|---|---|
| CF0134, CF0135, CF0517,CF0291, CF0304, CF0594 | F508del/711+1G>T |
| CF0138 | F508del/(TG)12(T)5 |
| CF0139 | 1677delTA/IVS16+1G>A(3120+1G>A) |
| CF0140 | S1251N/R117H |
| CF0141 | F508del/S945L |
| CF0160 | A455E/E60X |
| CF0167, CF0126, CF0196, CF0124, CF0099, CF0190, CF0154, CF0067, CF0109, CF0216, CF0061, CF0141, CF0053, CF0114, CF0112 | F508del/S1251N |
| CF0171 | F508del/Gly1349fs |
| CF0174 | 2105-2117del13insAGAAA/Q1352H |
| CF0175 | F508del/365-366insT(W79fs) |
| CF0181 | 1717-1G>A/2183AA>G |
| CF0191 | F508del/4374+1G>T |
| CF0198 | R553X/c.4375-3T>A |
| CF0217 | F508del/E730X |
| CF0219, CF0238, CF0176 | F508del/R347P |
| CF0224 | 711+1G>T/711+1G>T |
| CF0225 | F508del/L1034P |
| CF0227 | 3272-26A>G/G970R |
| CF0231, CF0228, CF0607, CF0606 | F508del/3272-26A>G |
| CF0236 | 711+1G>T/CFTRdele11 |
| CF0253 | F508del/Q1313X |
| CF0256 | R334W/R334W |
| CF0262 | F508del/D1152H |
| CF0269 | F508del/G85E |
| CF0270 | F508del/2184delA |
| CF0271, CF0168, CF0271, CF0188 | 1811+1G>C/1811+1G>C |
| CF0272 | E92K/E92K |
| CF0278 | F508del/W846X |
| CF0289 | R1162X/3659delC |
| CF0296 | N1303K/G550X |
| CF0297 | F508del/1294del7 |
| CF0303 | A455E/3659delC |
| CF0307 | F508del/3849+5G->A |
| CF0308 | F508del/621+1G>T |
| CF0317 | F508del/CFTRdele19.20 |
| CF0328 | F508del/2184insA |
| CF0334 | R1066H/CFTRdele2.3 |
| CF0335, CF0593 | F508del/1078delT |
| CF0340 | F508del/Y109D |
| CF0341 | F508del/S489X |

| | |
|---|---|
| CF0342, CF0375 | F508del/2143delT |
| CF0346 | F508del/1813insC |
| CF0348 | F508del/R74P |
| CF0349 | F508del/4382delA |
| CF0357 | R1162X/3539del16 |
| CF0358 | F508del/c.1725-1727del insAT |
| CF0359 | F508del/394delTT |
| CF0360, CF0361 | A46D/A46D |
| CF0373, CF0520 | F508del/Y849X |
| CF0378 | F508del/3905insT |
| CF0388 | F508del/G550X |
| CF0391 CF0822 | G542X/R1066C |
| CF0392 | W1282X/L927P |
| CF0396 | Y849X/2789+5G>A |
| CF0398 | 1717-1G>A/3905insT |
| CF0400 | R1066C/R1066H |
| CF0401 | 3849+10kbC>T/R347H |
| CF0406 | F508del/4383delA |
| CF0412 | Y275X/A559T |
| CF0414 | 1078delT/3272-26A>G |
| CF0419 | A455E/1343delG |
| CF0422, CF0458, CF0215 | F508del/G628R |
| CF0432, CF0403 | F508del/R1066C |
| CF0437 | F508del/IVS11-1G>C |
| CF0442 | F508del/4016insT |
| CF0460 | F508del/L732X |
| CF0477 | F508del/3659delC |
| CF0478 | 3905insT/D1152H |
| CF0480 | F508del/R75Q |
| CF0487 | F508del/L206W |
| CF0488 | F508del/Q493X |
| CF0489 | G542X/W679X |
| CF0523, CF0623, CF0326 | F508del/S18I |
| CF0534 | F508del/I1027T |
| CF0551 | F508del/1342-1delG |
| CF0556 | F508del/c.4243-3T>A |
| CF0563 | N1303K/c.3035A>C |
| CF0570 | (TG)12(T)7/(TG)11(T)7 |
| CF0572, CF0597 | R1162X/D1152H |
| CF0574 | F508del/G576A |
| CF0576 | 3272-26A>G/1898+5G>T |

| | |
|---|---|
| CF0579 | N1303K/G85E |
| CF0583 | F508del/1682dup |
| CF0588 | L732X/L732X |
| CF0592, CF0248, CF0384, CF0621, CF0433 | F508del/CFTRdele17a.17b |
| CF0596 | F508del/I507del |
| CF0600 | F508del/R1158X |
| CF0608 | R785X/R785X |
| CF0610 | F508del/R1358S |
| CF0612 | 3272-26A>G/3272-26A>G |
| CF0641 | G85E/1677delTA |
| CF0645 | R1162X/3849+10kbC>T |
| CF0665 | F508del/I336K |
| CF0667 | F508del/S586N |
| CF0669 | L1335P/L1335P |
| CF0671 | F508del/H620P |
| CF0696 | R553X/4005+2T-c |
| CF0699 | F508del/L453S |
| CF0705 | F508del/T1396P |
| CF0706, CF0187 | F508del/(TG)11(T)5 |
| CF0710 | S1251N/1717-1G>A |
| CF0712 | F508del/G1249E |
| CF0715 | 4382delA/2043delG |
| CF0733 | L206W/S1235R |
| CF0744 | V1160T/E92K |
| CF0754 | F508del/Ile336fs |
| CF0823 | G542X/P988R |
| CF0829 | A455E/711+5G>T |
| CF0841 | F508del/R352W |
| CF0844 | F508del/c.3407_3422del |

**Supplemental Table 5:** Overview of genotypes and corresponding mutation category of reference PDIOs included in CFTR modulator screen, corresponding to **Figure 5C & 5D**

| Category | Genotype | N |
|---|---|---|
| F508del/RF_splice | F508del/2789+5G>A | 3 |
| | F508del/3272-26A>G | 4 |
| | F508del/3849+10kbC>T | 5 |
| F508del/RF_missense | F508del/G551D | 2 |
| | F508del/D1152H | 1 |
| | F508del/L206W | 1 |
| | F508del/S945L | 1 |
| | F508del/A445E | 4 |
| S1251N/other | S1251N/F508del | 15 |
| | S1251N/1717-1G>A | 1 |
| R117H/other | R117H/F508del | 6 |
| | R117H/R1162X | 1 |
| | R117G/R553X | 1 |
| | R117H/T1857delT | 1 |
| | R117H/W1282X | 1 |

| | | |
|---|---|---|
| F508del/minimal | F508del/W1282X | 5 |
| | F508del/R1066C | 2 |
| | F508del/Y1092X | 5 |
| | F508del/N1303K | 5 |
| | F508del/G85E | 2 |
| | F508del/1078delT | 1 |
| | F508del/1717-1G>A | 4 |
| | F508del/2184delA | 2 |
| | F508del/I507del | 1 |
| | F508del/R553X | 1 |
| | F508del/1078delT | 1 |
| | F508del/2143delT | 2 |
| | F508del/2183AA>G | 1 |
| | F508del/3659delC | 2 |
| | F508del/3905insT | 1 |
| | F508del/394delTT | 1 |
| | F508del/CFTRdele2.3 | 3 |
| | F508del/G542X | 1 |
| F508del/F508del | F508del/F508del | 1 |
